# Beta-frequency sensory stimulation enhances gait rhythmicity through strengthened coupling between striatal networks and stepping movement

**DOI:** 10.1101/2024.07.07.602408

**Authors:** Sudiksha Sridhar, Eric Lowet, Howard J. Gritton, Jennifer Freire, Chengqian Zhou, Florence Liang, Xue Han

**Affiliations:** Department of Biomedical Engineering, Boston University, Boston, MA, USA; Department of Neuroscience, Erasmus MC, Rotterdam, the Netherlands; Department of Comparative Biosciences, University of Illinois at Urbana-Champaign, Urbana, IL, USA; Department of Pharmacology, Boston University, Boston, MA, USA

## Abstract

Stepping movement is delta (1-4 Hz) rhythmic and depends on sensory inputs. In addition to delta rhythms, beta (10-30 Hz) frequency dynamics are also prominent in the motor circuits and are coupled to neuronal delta rhythms both at the network and the cellular levels. Since beta rhythms are broadly supported by cortical and subcortical sensorimotor circuits, we explore how beta-frequency sensory stimulation influences delta-rhythmic stepping movement, and dorsal striatal circuit regulation of stepping. We delivered audiovisual stimulation at 10 Hz or 145 Hz to mice voluntarily locomoting, while simultaneously recording stepping movement, striatal cellular calcium dynamics and local field potentials (LFPs). We found that 10 Hz, but not 145 Hz stimulation prominently entrained striatal LFPs. Even though sensory stimulation at both frequencies promoted locomotion and desynchronized striatal network, only 10 Hz stimulation enhanced the delta rhythmicity of stepping movement and strengthened the coupling between stepping and striatal LFP delta and beta oscillations. These results demonstrate that higher frequency sensory stimulation can modulate lower frequency dorsal striatal neural dynamics and improve stepping rhythmicity, highlighting the translational potential of non-invasive beta-frequency sensory stimulation for improving gait.

## Introduction

Stepping movement is highly rhythmic and occurs at delta frequencies (1-4 Hz) in humans and mice^1,2^. While the spinal central pattern generators are critical for pacing movement^3,4^, stepping related delta-rhythmic neural activities are broadly detected in many motor-related brain areas, including the motor cortex^5–7^, dorsal striatum^8^, subthalamic nucleus (STN)^9,10^ and cerebellum^7,11,12^. Stepping, as with many aspects of voluntary movement, is intricately linked to sensory inputs^13^ and the reliance on sensory cues is often used as a compensation strategy to improve gait in Parkinsonian patients^14–16^. Indeed, rhythmic auditory cues, such as metronome, music with adapted tempo or beats, or counting, when played around individual subject’s stepping frequencies have been shown to improve patients’ gait and mobility^17–22^.

Beta frequency (8-35 Hz) oscillations are broadly observed across the cortical-basal ganglia circuits and related to sensorimotor processing^23–27^. During locomotion, transient movement related LFP beta bursts, hundreds of milliseconds long, are prominent across the motor cortex, striatum, globus pallidus externus, and subthalamic nucleus (STN) ^9,23^. In the striatum, transient LFP beta oscillations have been reported during task initiation^26^, cue processing^23^ and movement completion^24^. However, persistent, and exaggerated beta oscillations are widely recognized as a functional biomarker of akinesia and bradykinesia in Parkinson’s disease^28,29^. Thus, beta oscillations are hypothesized to maintain status quo^25,27^, as transient beta also occurs during motor planning without actual movement, and inhibition or desynchronization of beta promotes movement transitions. Consistent with the anti-kinetic effects of persistent beta rhythms, transcranial alternating current stimulation (tACS) at beta frequencies inhibits motor learning^30^, slows voluntary movement, and reduces force generation in humans^31,32^. Similarly, beta-frequency deep brain stimulation (DBS) in the STN leads to further deterioration of bradykinesia and rigidity in Parkinsonian patients^33–35^.

Dorsal striatum, the largest input structure of the basal ganglia, receives sensory and motor inputs from broad cortical areas^36,37^ including somatosensory, visual, auditory and motor cortices, and from many thalamic areas^38,39^ such as centromedian, parafascicular, centrolateral, pulvinar and lateral posterior/dorsal nuclei. Striatum plays important roles in various aspects of motor control and motor learning^40^ and is responsive to sensory stimulation^41^. Individual striatal neurons exhibit diverse responses during sensorimotor tasks and naturalistic behaviors^42^. At the network level, recent large-scale cellular calcium imaging analyses revealed that functional connectivity between clusters of striatal neurons could increase during specific actions^43,44^, even though the overall network correlation strength may decrease^45^, highlighting dynamic recruitment of subpopulations of striatal neurons during behaviors.

The coupling of delta and beta LFP oscillations are broadly observed across the cortico-basal ganglia-thalamic circuits^2,28^. Specifically, fluctuation of striatal beta power was found to be coupled to the phase of cortical and thalamic neuronal delta oscillations^2,28,46^. Intriguingly, striatal beta was also temporally locked to internally generated delta rhythmic finger tapping behavior^47^, suggesting that both neuronal and behavioral delta rhythms can temporally organize beta rhythmicity^48,49^. Recently, using cellular voltage imaging, we discovered that the membrane potentials of many striatal neurons exhibit prominent delta oscillations, which organize beta-rhythmic spike bursting and striatal LFP beta oscillations^8^, highlighting a potential cellular mechanism for the observed circuit level beta and delta coupling.

While strong stimulation of the motor circuits at beta frequencies using tACS or DBS results in anti-kinetic effects^30–35^, we hypothesized that sensory stimulation at beta frequencies could boost motor circuit processing of stepping movement via sensory entrainment, without producing exaggerated beta oscillations that are known to be anti-kinetic. To test this hypothesis, we delivered beta rhythmic audiovisual stimulation to mice voluntarily locomoting, since low frequency stimulations, sensory, tACS, DBS, and transcranial magnetic stimulation (TMS) have been found to entrain endogenous neural circuit oscillations^50–53^. The exact beta bands vary widely across animal models, behavioral states and individual Parkinsonian patients^54^. While beta frequencies generally cover ∼12-30 Hz, they can span 8-35 Hz^55–58^. In awake, head-fixed mice, our previous studies found that striatal beta band centers around 10 Hz^45,55^, and thus we delivered stimulation at 10 Hz to maximize the chance of entraining cortical-striatal circuits. We compared the effect of 10 Hz sensory stimulation to the higher frequency 145 Hz stimulation that would be less effective in entraining neural circuits^53^. We examined the relationships between cellular calcium dynamics, functional connectivity between simultaneously recorded neurons, population striatal LFPs, and stepping movement.

## Results

### Population striatal LFP dynamics were better entrained by sensory stimulation at 10 Hz than 145 Hz

To examine how non-invasive sensory stimulation at beta frequencies influences basal ganglia circuitry and locomotion, we performed calcium imaging from individual striatal neurons in head fixed mice freely navigating on a spherical treadmill (**Fig. 1a**). Briefly, striatal neurons were transduced with the genetically encoded calcium indicator GCaMP7f through intracranial infusion of AAV9-syn-GCaMP7f (n=5 mice) or AAV9-syn-SomaGCaMP7f (n=4 mice). GCaMP7f expressing neurons were then recorded using a custom microscope through an imaging window positioned above the dorsal striatum that included the conventionally defined dorsolateral and dorsomedial striatum (**Fig 1a**). We also recorded LFPs through a nearby electrode coupled to the imaging window, about 200 µm away from the imaging site (**Fig. 1a**). During each experiment, mice were presented with five one-minute-long audiovisual stimulation every five minutes, resulting in a total recording of 36 minutes per session (**Fig. 1b**). The audiovisual stimulation consisted of concomitant auditory clicks (∼77 dB, 50% duty cycle) and visual flashes (50% duty cycle) pulsed at 10 Hz or 145 Hz, in a dimly lit room with ∼72 dB constant background noise.

**Figure 1:**
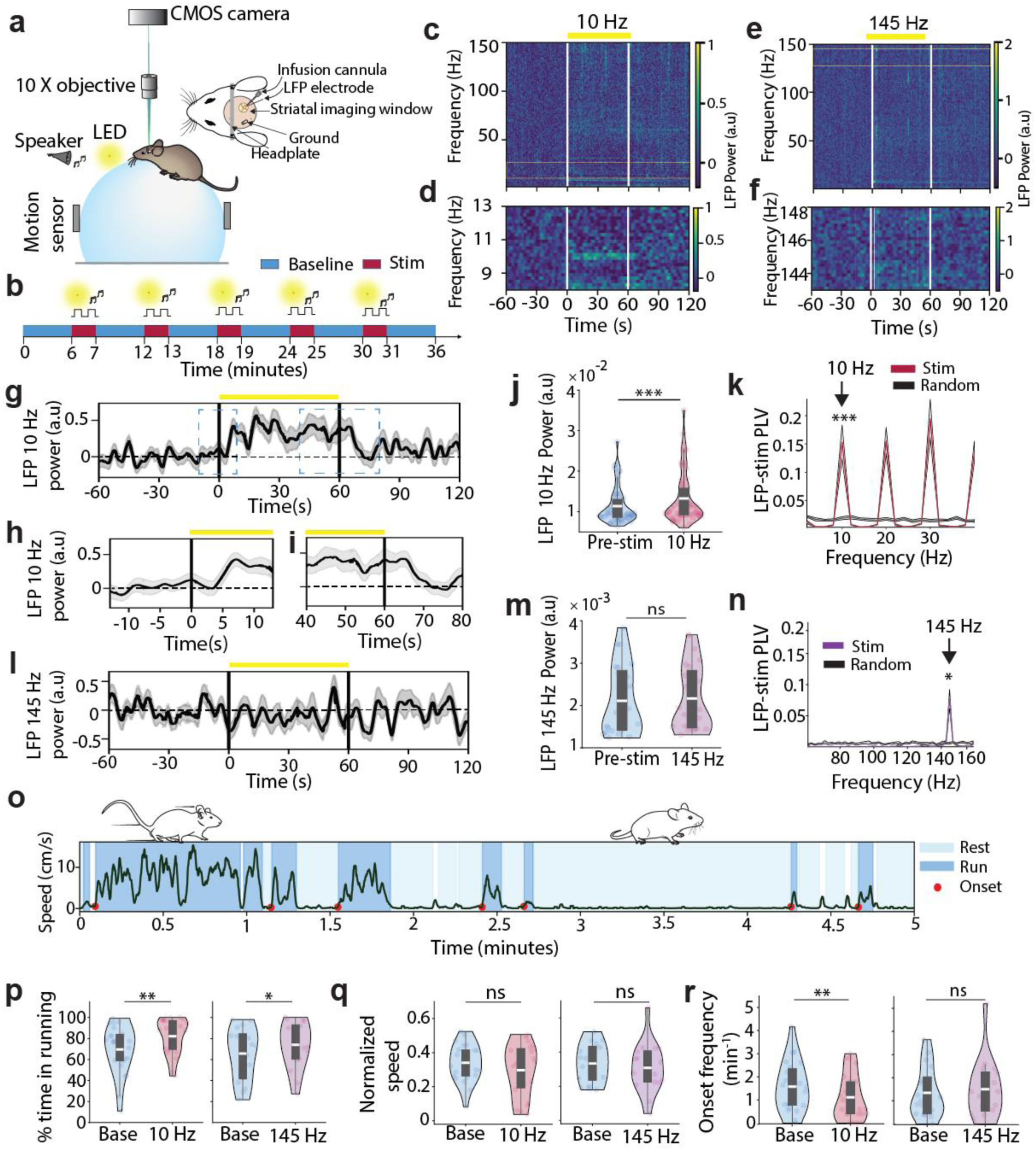
10 Hz sensory stimulation prominently entrained striatal LFPs and suppressed movement onset transitions. **(a)** Illustration of experimental setup with a head-fixed mouse under a custom microscope voluntarily locomoting on a treadmill. **(b)** Experimental timeline. **(c)** Normalized LFP power spectrum aligned to stimulation onset during 10 Hz stimulation. LFP power was Z-scored to the mean pre-stimulation period. **(d)** Zoom in around 10 Hz as highlighted by the box in (c). **(e, f)** Same as (**c, d**) but during 145 Hz stimulation. **(g)** Mean 10 Hz LFP power aligned to stimulation onset across all 10 Hz stimulation sessions. **(h, i)** Zoom in around 10 Hz stimulation onset (**h**) and offset (**i**). **(j)** 10 Hz power during 10 Hz stimulation vs. pre-stimulation period (Wilcoxon signed rank test, p=8.9e^-5^, n=65 trials from 13 recording sessions). **(k)** Mean phase-locking value (PLV) between 10 Hz stimulation pulse trains and LFP across frequencies during stimulation (red) vs. random shuffles (black). PLV at 10 Hz is significantly greater during stimulation than random shuffles (Wilcoxon signed rank test, p=2.4e^-4^, n=65 trials). Shaded area is standard error of mean. **(l)** Mean 145 Hz LFP power aligned to stimulation onset across all 145 Hz stimulation sessions. **(m)** 145 Hz power during 145 Hz stimulation vs. pre-stimulation period (Wilcoxon signed rank test, p=0.44, n=24 trials). **(n)** Mean PLV of 145 Hz stimulation pulse train to LFP during stimulation (purple) vs. random shuffles (black) (Wilcoxon signed rank test, p= 0.016). **(o)** An example locomotion speed recording, showing running bouts (dark blue shading), resting periods (light blue shading), and onset transitions (red stars). **(p)** The percentage of time mice spent on running during 10 Hz or 145 Hz stimulation vs. baseline. Stimulation increased running time (Wilcoxon signed rank test, 10 Hz: p=0.0014, n=23 sessions; 145 Hz: p=0.022, n=17 sessions). **(q)** Normalized speed (division by maximum speed of a session) during running, upon 10 Hz or 145 Hz stimulation relative to baseline. There was no difference (Wilcoxon signed rank test, 10 Hz: p=0.19, n=23 sessions; 145 Hz: p=0.98, n=17 sessions). **(r)** Onset transition frequencies during 10 Hz and 145 Hz stimulation relative to baseline. Onset transition is significantly reduced for 10 Hz, but not 145 Hz stimulation (Wilcoxon signed rank test, 10 Hz: p=0.004, n=23 sessions; 145 Hz: p=0.523, n=17 sessions). Quantifications are visualized as violin plots with the outer shape representing the data kernel density, and a box plot (box: interquartile range, whiskers: 1.5x interquartile range, white line: mean). *p < 0.05, **p < 0.01, ***p < 0.001

We first evaluated the sensory entrainment of LFPs by comparing LFP spectral power during stimulation versus the one-minute pre-stimulation period. As expected, 10 Hz stimulation significantly increased 10 Hz LFP power during stimulation (**Fig. 1c-d, 1g, 1j**). Further, the population LFP power increase at 10 Hz lagged the stimulation onset by 5.5 sec (**Fig 1g-h)** and remained elevated for 8.5 sec after stimulation offset (**Fig. 1g, 1i**). Finally, 10 Hz stimulation exhibited significant phase locking to the 10 Hz component of the LFP signals (**Fig. 1k**), during both running and resting (**Supplementary Fig. 1a, b**), further confirming the entrainment effect is independent of movement states.

In contrast, 145 Hz stimulation did not alter the 145 Hz LFP spectral power (**Fig. 1e-f**, **l-m**). Interestingly, we detected a weak but nonetheless significant phase locking between 145 Hz stimulation and the 145 Hz component of the LFPs, though the phase locking strength was an order of magnitude weaker than that observed with 10 Hz stimulation (**Supplementary Fig. 1c**, **Fig. 1n**). Moreover, 10 Hz stimulation did not alter 145 Hz LFP power (**Supplementary Fig. 1d)**, and 145 Hz stimulation did not alter 10 Hz LFP power (**Supplementary Fig 1e)**, suggesting that the observed entrained effects were specific to the frequencies of audiovisual stimulation. Together, these results demonstrate that audiovisual stimulation at 10 Hz produces prominent entrainment of the striatal network, whereas stimulation at 145 Hz mediates a much weaker effect.

### Sensory stimulation at either 10 Hz or 145 Hz increased the time mice spent on running, but only 10 Hz suppressed movement onset transitions

We next examined the influence of sensory stimulation on locomotion. Since each recording session consisted of five stimulation trials at either 10 Hz or 145 Hz, movement features were compared between stimulation versus baseline of the same recording session. Baseline was the period excluding the stimulation periods and the minute after each stimulation to avoid any residual stimulation effects (**Fig. 1o**). We first identified running bouts defined as continuous movement for at least 2 seconds, and the corresponding onset transitions. We found that sensory stimulation at both frequencies increased the fraction of time mice spent on running (**Fig. 1p, Supplementary Fig. 2a**), without changing the mean velocity during running (**Fig. 1q**). Interestingly, 10 Hz stimulation, but not 145 Hz stimulation, led to a significant reduction in movement onset transitions and longer running bout durations (**Fig. 1r**, **Supplementary Fig. 2c-d)**, consistent with the general notion that beta rhythms help maintain the status quo^25,27^.

Striatal LFPs are known to be modulated by locomotion^59^. During baseline without stimulation, we found that LFP theta (6-8 Hz) and gamma (45-80 Hz) power increased during movement (**Supplementary Fig. 1f-g**). The theta power increase was particularly prominent with a peak at ∼7 Hz and a harmonic at ∼14 Hz. However, the narrow band 10 Hz beta power was suppressed during running in the absence of stimulation (**Supplementary Fig. 1g**). Due to the strong theta harmonic, we focused on higher frequency beta (15-30 Hz) band, which was also decreased during running (**Supplementary Fig. 1f-g)**, consistent with the general suppression of beta power during movement^25^ and our previous findings in mice under similar behavioral conditions^45^. While sensory stimulation at either 10 Hz or 145 Hz did not change the LFP power at theta, beta, and gamma bands during rest (**Supplementary Fig. 1h-i)**, 10 Hz stimulation suppressed the beta power even further than baseline without stimulation, whereas theta power increased compared to the baseline (**Supplementary Fig. 1j**, l**-m**). Together, these results demonstrate that even though sensory stimulation at both frequencies promoted movement, only 10 Hz stimulation, but not 145 Hz, suppressed movement onset transitions. These behavioral results along with the differential effects of the two stimulation frequencies on striatal LFPs highlight that beta-frequency sensory stimulation is effective at reducing movement flexibility, likely through entrainment of locomotor circuits.

### 10 Hz stimulation, but not 145 Hz, enhanced stepping rhythmicity and improved the coupling between stepping and LFP beta

Since locomotion in mice and humans occurs in the delta frequency range^1^, we calculated the spectral power of the treadmill speed, as treadmill speed fluctuations closely track animals stepping cycles during running^8,60^. Consistent with our previous observations under similar locomotion conditions^8^, mice stepping occurred at delta frequencies with a peak around 3-4 Hz (**Fig. 2a-b**). Upon 10 Hz stimulation, animal’s movement speed during running became more rhythmic around delta frequencies (3-4 Hz), showing a significant increase in delta power (**Fig. 2c, 2e**). In contrast, during 145 Hz stimulation, the power spectral density of movement speed exhibited a much wider range, and 145 Hz stimulation did not alter movement speed delta power (**Fig. 2d-e**). After detecting a change in stepping rhythmicity during 10 Hz stimulation, we next examined how sensory stimulation influences the temporal relationship between striatal population LFP dynamics and stepping by computing the phase-locking value (PLV) between LFPs and movement speed. During running, in the absence of sensory stimulation, there was a significant phase locking between LFPs and movement speed at delta frequencies (**Fig. 2f-h**), consistent with our previous findings^8^. Interestingly, 10 Hz stimulation, but not 145 Hz stimulation, further promoted LFP-movement speed phase locking at delta frequencies during running (**Fig. 2f-h**).

**Figure 2:**
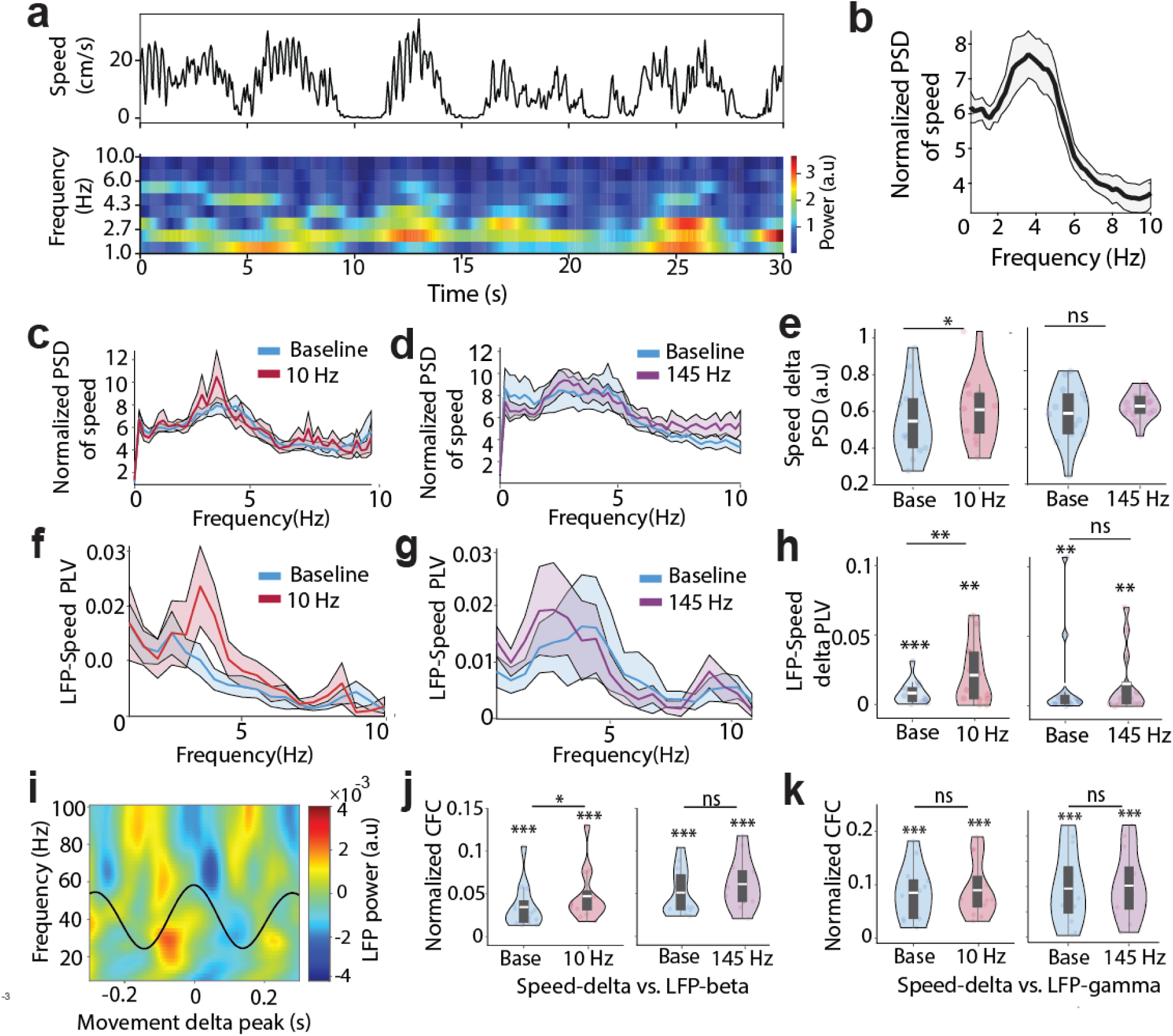
10 Hz, but not 145 Hz, enhanced gait rhythmicity and elevated coupling of stepping with striatal LFP delta and beta oscillations. **(a)** Example movement speed trace (top) and corresponding power spectrogram (bottom) showing delta rhythms. **(b)** Power spectral density (PSD) of movement speed during running normalized to resting across all recording periods. Black line corresponds to the mean across all recording sessions, and shading, the standard error of the mean (n=28 recording sessions). **(c)** Similar to (**b**), but for 10 Hz stimulation sessions, with baseline in blue and 10 Hz stimulation in red. **(d)** same as (**c**), but for 145 Hz stimulation sessions. **(e)** Quantification of speed delta (3-4 Hz) PSD across all running bouts during baseline vs. stimulation (Wilcoxon signed rank test, 10 Hz, p=0.01, n=14 sessions; 145 Hz: p=0.17, n=14 sessions). **(f)** LFP-movement speed PLV across frequencies during baseline (blue) vs. 10 Hz stimulation (red). **(g)** same as (f), but for 145 Hz stimulation. **(h)** LFP-movement speed delta PLV during baseline vs. stimulation (Wilcoxon signed rank test, 10 Hz: baseline vs. shuffle, p=2.4e^-4^, stimulation vs. shuffle: p=0.001, baseline vs. stimulation: p=0.008, n=13; 145 Hz: baseline vs. shuffle: p=0.002, stimulation vs. shuffle: p=0.006, baseline vs. stimulation: p=0.74, n=13). **(i)** Heatmap showing the relative LFP power aligned to the peak of the delta-frequency component of movement speed. **(j)** Cross frequency coupling (CFC) between movement speed delta and beta-filtered LFP during baseline vs. stimulation (Wilcoxon signed rank test, 10 Hz: baseline: p=2.4e^-4^, 10 Hz: p=2.4e^-4^, baseline vs 10 Hz: p=0.03; 145 Hz: baseline: p=2.4e^-4^, 145 Hz: p=2.4e^-4^, baseline vs 145 Hz: p=0.24). **(k)** CFC between movement delta and gamma-filtered LFP during baseline vs. stimulation (Wilcon signed rank test, 10 Hz: baseline: p=2.4e^-4^, 10 Hz: p=2.4e^-4^, baseline vs. 10 Hz: p=0.45; 145 Hz: baseline: p= 2.4e^-4^, 145 Hz: p=2.4e^-4^, baseline vs. 145 Hz: p=0.73). Lines represent the mean across recording sessions and shaded regions are the standard error of the mean. Quantifications are visualized as violin plots with the outer shape representing the data kernel density and a box plot (box: interquartile range, whiskers: 1.5x interquartile range, white line: mean). *p < 0.05, **p < 0.01, ***p < 0.001.

Our previous study also showed that striatal LFP beta and gamma power are nested within the delta-rhythmic stepping cycles, peaking at a similar phase as spikes in delta-rhythmic striatal neurons^8^. To understand how sensory stimulation influences the relationship between higher frequency components of the LFP power and stepping movement, we filtered the movement speed trace in the broader delta frequency range (2-4 Hz) and computed the mean LFP spectrogram around all movement delta-peaks. Consistent with our previous observations^8^, in the absence of stimulation, LFP beta and gamma power (40-100 Hz) were modulated by the movement delta (**Fig. 2i-k**).

During stimulation at both 10 Hz and 145 Hz, the nesting of LFP beta and gamma within the delta-rhythmic movement speed trace remained evident (**Fig. 2j-k**). Thus, delta-rhythmic movement organizes the timing of LFP beta and gamma regardless of sensory stimulation. Interestingly, 10 Hz stimulation, but not 145 Hz, selectively increased the coupling between movement delta phase and LFP beta power, but not gamma power (**Fig. 2j, k**). Together, these results demonstrated that while audiovisual stimulation at both 10 Hz and 145 Hz promoted movement (**Fig. 1p**), only 10 Hz stimulation enhanced the delta rhythmicity of stepping (**Fig. 2c, e**), which was accompanied by strengthened delta-frequency phase locking between LFP and movement and increased coupling between LFP beta power and movement delta phase (**Fig. 2j**). These results highlight beta-frequency specific sensory modulation of striatal involvement in coordinating stepping on a moment-to-moment basis through improved coupling of striatal neural activity and movement.

### Cellular calcium activity preceded locomotion

To examine how sensory stimulation modulates striatal encoding of locomotion, we performed single cell calcium imaging from hundreds of striatal neurons simultaneously using GCaMP7f that is better at capturing spiking than the older generations of GCaMPs^44^ (**Fig. 3a-b**). While intracellular calcium concentration fluctuates due to various biochemical and biophysical activities, the rising phase of the transiently occurring calcium events is generally associated with elevated spiking probabilities, whereas the falling phase is influenced by the slow kinetics of calcium ion dissociation from GCaMP7f and calcium clearance from the cytosol. Thus, we identified calcium events in the GCaMP7f fluorescence traces and binarized the event rising phase as ones and everywhere else as zeros to capture periods of heightened neuronal activity (**Supplementary Fig. 3a**). During baseline without stimulation, we detected 2.45 ± 1.58 calcium events /min (mean ± standard deviation, n=6409 neurons from 24 sessions), consistent with previous calcium imaging studies of striatal neurons using GCaMP7f^44^, but higher than that using GCaMP6f^43,45^. Using the rising phase of each calcium event, we obtained an activity rate of 0.24 ± 0.15 Hz (mean ± standard deviation, n=6096 neurons from 24 sessions). As with our previous study^44^, we detected no difference between GCaMP7f and Soma-GCaMP7f in terms of calcium event rate, activity rate, or pairwise Pearson correlation coefficients (**Supplementary Fig. 3b-d**). Thus, all subsequent analyses were performed by combining the two mice groups.

**Figure 3:**
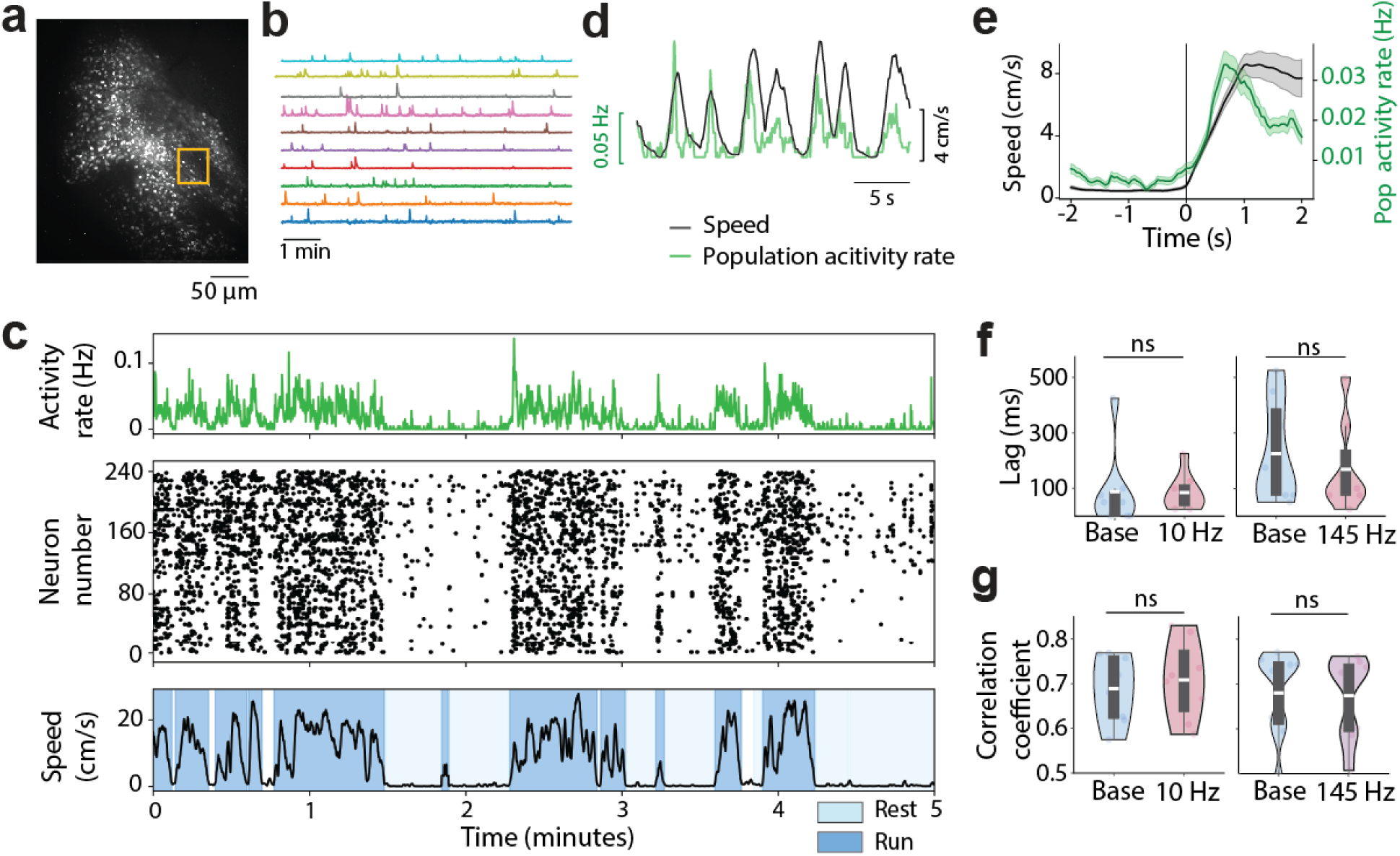
Striatal calcium activity preceded locomotion. **(a)** A maximum-minimum GCaMP7 fluorescence intensity image from an example recording session. **(b)** Example fluorescence traces from neurons in the yellow box in (**a**). **(c)** An example recording showing the mean calcium activity rate across movement-responsive neurons (top), the raster plot displaying calcium activity rate across individual neurons (middle), and the corresponding movement speed (bottom), with resting (light blue) and running (dark blue) bouts highlighted. **(d)** Instantaneous population activity rate (green) across movement-responsive neurons during one example recording session and the corresponding movement speed (black). **(e)** Average population activity rate of movement-responsive neurons (green) in the same example session as in (d) aligned to movement onsets (average speed trace in black). **(f)** The time lags between activity rate and movement speed during baseline vs. stimulation (Wilcoxon signed rank test, 10 Hz: p=0.46, n=8; 145 Hz, p=0.16, n=8). **(g)** The peak correlation coefficients during baseline vs. stimulation (Wilcoxon signed rank test, 10 Hz, p=0.5, n=8; 145 Hz, p=0.58, n=8). Quantifications are visualized as violin plots with the outer shape representing the data kernel density and a box plot (box: interquartile range, whiskers: 1.5x interquartile range, white line: mean). *p < 0.05, **p < 0.01, ***p < 0.001.

As many striatal neurons are known to increase their activity during movement^43,45,61^, we first determined the fraction of neurons that were modulated by locomotion under our experimental conditions. Briefly, we computed the difference in activity rate between running and resting and then compared it to a shuffled distribution computed from two randomly chosen baseline periods (details see **Methods**). Since calcium activity rate is generally low, we identified movement-responsive neurons as those with a significant increase in activity rate (>97.5^th^ percentile in shuffled distribution) during running compared to resting. During the baseline period without sensory stimulation, we found 70.4% neurons were activated by movement (**Fig. 3c**, 4289/6096 neurons from 24 sessions), with a mean activity rate of 0.44 ± 0.24 Hz during movement across movement-responsive neurons.

We also noticed that the activity of movement-responsive neurons preceded the rise in speed (**Fig. 3d-e**). To quantify this temporal relationship, we computed the cross correlation between the population activity of movement-responsive neurons and locomotion speed with different temporal lags and identified the time lag when the peak correlation occurred. Across baseline sessions without stimulation, the peak correlation was 0.68 ± 0.02 (mean ± standard error, n=16 sessions, **Supplementary Fig. 3e)**, with neuronal responses preceding locomotion by 156 ± 42.13 ms (mean ± standard error, n =16 sessions; range: 50-525 ms; median: 75 ms). This result is in general agreement with prior electrophysiology findings showing striatal neuron activation before movement^62,63^. Further, audiovisual stimulation did not alter the time lags (**Fig. 3f**) or the peak correlation coefficient (**Fig. 3g**). Thus, even though sensory stimulation heightened locomotion bouts (**Fig. 1p**), it did not disrupt the temporal relationship of striatal activity and locomotion.

### Cellular neuronal activity tracked delta-rhythmic movement and sensory stimulation did not alter this temporal relationship

After observing a temporal relationship between population LFP delta and movement stepping cycles, we further evaluated the relationship between individual neurons and stepping. As our imaging frame rate was 20 Hz, though sufficient to capture intrinsically slow cytosolic calcium dynamics, it limited our analysis to frequencies below 10 Hz. Surprisingly, we found that calcium event onsets exhibited significant phase locking to the broader delta (2-4 Hz) frequency component of the movement speed (**Fig. 4a-g**). Thus, cellular calcium dynamics tracks gait cycles, similar to previous studies that recorded striatal neuron spiking^8,64,65^. However, sensory stimulation at either frequency did not change the phase locking strength (**Fig. 4l, Supplementary Fig. 4a, c**), or the preferred phase (**Supplementary Fig. 4b, d**). Furthermore, event onsets were also phase locked to the delta component of the LFP trace (**Fig. 4a-c, 4h-k**), and stimulation did not alter the phase locking strength or the preferred phase (**Fig. 4m, Supplementary Fig. 4e-h**). The prominent phase locking of calcium events to stepping movement and LFP delta oscillations confirms that individual striatal neurons are coordinated with step-by-step movement, as demonstrated in previous studies^8,64,65^. The fact that sensory stimulation did not alter the temporal relationship between striatal cellular dynamics and stepping movement further support the notion that the physiological audiovisual stimulation boosts intrinsic striatal circuit processing of locomotion.

**Figure 4:**
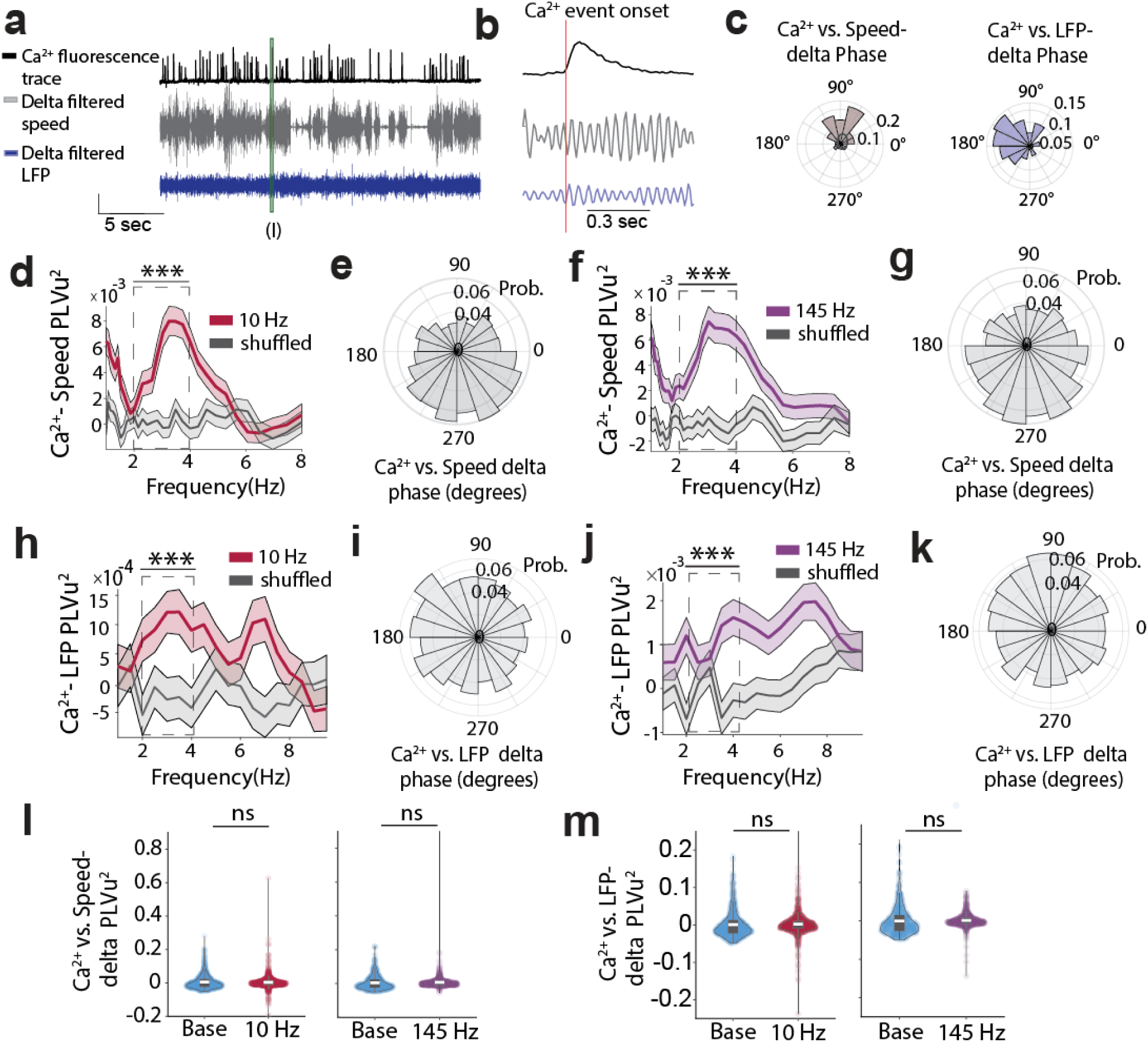
Striatal cellular activity was phase coupled to stepping movement and LFP delta oscillations and this temporal relationship was unaffected by sensory stimulation. **(a)** An example neuron’s fluorescence trace with the corresponding delta-frequency filtered movement speed and LFP traces. (**b**) Zoom-in view around a calcium event onset indicated by the green line in (a). (**c**) An example neuron’s polar histograms of the preferred phase of calcium events relative to movement speed delta (left) and LFP delta (right). Shown is the same neuron as in (a). **(d)** Population PLVu^2^ of calcium event onsets across all neurons to movement speed during running (red) and random shuffles (gray) throughout the entire recording session with 10 Hz stimulation. **(e)** Population polar histogram of calcium event phase to speed delta (2-4 Hz) across all neurons during 10 Hz stimulation sessions (Mixed effect models, 10 Hz: p=7.1e^-14^, n=2510). **(f,g)** Same as (d,e), but during 145 Hz stimulation sessions (Mixed effect models, p=1.2e^-10^, n=2669). **(h)** PLVu^2^ of calcium event onsets to LFP across frequencies during running (red) and random shuffles (gray) throughout the entire recording session with 10 Hz stimulation. There is significant phase locking at delta frequencies (Mixed effect models, p= 1.2e^-6^, n=2510). **(i)** Population polar histogram of calcium event onsets to LFP delta (2-4 Hz) across all neurons during 10 Hz stimulation sessions. **(j,k)** Same as (h,i), but during 145 Hz stimulation sessions (Mixed effect models, p=0.0025, n=2669)**. (l)** Quantification of Ca^2+^-speed delta PLVu^2^ during baseline vs. stimulation sessions (Mixed effect models, 10 Hz: p=0.27, n=907; 145 Hz: p=0.25, n=659). **(m)** Quantification of PLVu^2^ between calcium event onsets and LFP delta in baseline vs. stimulation (Mixed effect models, 10 Hz: p=0.44, n=889; 145 Hz: p=0.32, n=676). Lines represent the mean and the shaded regions represent the standard error of the mean. Quantifications are visualized as violin plots with the outer shape representing the data kernel density and a box plot (box: interquartile range, whiskers: 1.5x interquartile range, white line: mean). *p < 0.05, **p < 0.01, ***p < 0.001.

### Sensory stimulation at 10 Hz and 145 Hz modulated a greater and yet balanced fraction of activated versus suppressed neurons during movement

Aligning the response of individual neurons to audiovisual stimulation onset, we found that stimulation evoked responses were most prominent at the stimulation onset, with some neurons exhibiting increased calcium activity rates and others decreased rates (**Fig. 5a, 5d**). As we did not detect any trend across the five stimulation trials (**Supplementary Fig. 5a-f)**, evoked responses were combined across trials. As a population, evoked responses showed a sharp increase at stimulation onset, peaking around 0.3 s after onset, and then settling into a sustained increase after about 1 second, for both stimulation frequencies (**Fig. 5b, 5e**). While the transient population increase within one second of stimulation onset was significant for both frequencies (**Fig. 5g**), the sustained increase was only significant for 145 Hz stimulation, but not for 10 Hz (**Fig. 5h**). At stimulation offset, we also detected a significant rebound, peaking around 0.5 s after the offset (**Fig. 5c, 5f, 5i**).

**Figure 5:**
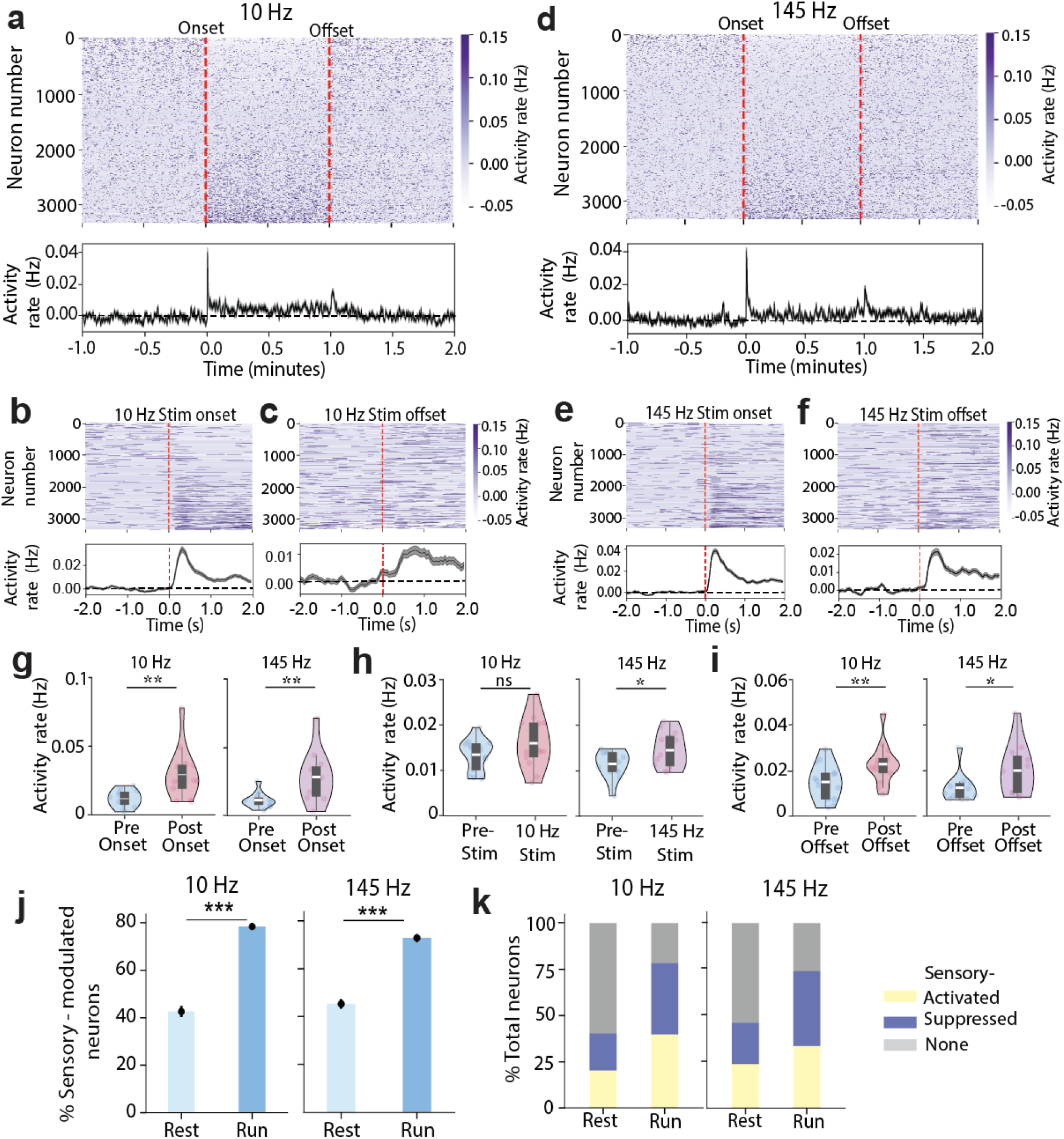
Sensory stimulation at 10 Hz and 145 Hz evoked a transient increase in activity rate at both onset and offset, and about equal fractions of neurons were activated and suppressed. (**a)** Heatmap of the activity rate of all neurons before, during and after 10 Hz stimulation (top), and their population average (bottom). Red dotted lines correspond to stimulation onset and offset. **(b, c)** Zoom-in view around the onset (b) and offset (c). **(d-f)** same as (**a**) but for 145 Hz stimulation. **(g)** Quantification of transient activity rate during 1 second period pre-vs. post-onset of stimulation (Wilcoxon signed rank test, 10 Hz: p=0.003, n=14; 145 Hz, p=0.005, n=12). **(h)** Quantification of sustained activity rate during 1 minute pre-vs. post-onset of stimulation (Wilcoxon signed rank test, 10 Hz: p=0.54, 145 Hz: p=0.016). **(i)** Quantification of the transient activity rate during 1 second pre-vs. post-offset of stimulation (Wilcoxon signed rank test, 10 Hz: p=0.001; 145 Hz: p=0.02). **(j)** Quantification of the fraction of modulated neurons during resting (light blue) vs. running (dark blue) for 10 Hz and 145 Hz sessions (Fisher’s test: 10 Hz: p=1.17e^-142^; 145 Hz: p=5.24e^-89^). **(k)** Stacked bar plot visualization of the proportions of sensory-activated (yellow), sensory-suppressed (indigo) and non-modulated neurons (gray) in each movement condition. Quantifications are visualized as violin plots with the outer shape representing the data kernel density and a box plot (box: interquartile range, whiskers: 1.5x interquartile range, white line: mean). *p < 0.05, **p < 0.01, ***p < 0.001.

Next, we determined whether each neuron was modulated, either positively or negatively, by sensory stimulation by comparing the activity rate during resting or running separately using the shuffling procedure as that for determining movement-responsive neurons (details in Methods).

During resting, about 44% neurons were modulated by sensory stimulation (**Fig. 5j**, **Table 1**), and during running, a significantly larger fraction (∼75.5%) were modulated (**Fig. 5j**, **Table 1-2,** Fisher’s test, 10 Hz: p=1.17e^-142^; 145 Hz: p=5.24e^-89^). Intriguingly, the proportions of positively and negatively modulated neurons were similar during both resting and running for both stimulation frequencies (**Fig. 5k**, **Table 1**).

### Sensory stimulation inhibited movement-responsive neurons during running

Further evaluation of movement-responsive neurons revealed that ∼84% were also responsive to sensory stimulation (**Fig. 6a**, **Table 3),** suggesting that many striatal neurons integrate sensory and motor information. We next characterized how sensory stimulation modulates movement-responsive neurons and found that more neurons were inhibited by sensory stimulation during running than during resting (**Fig. 6b**, **Supplementary Fig. 5g**, **Table 4**, Fisher’s test, 10 Hz: p=3.4e^-64^; 145 Hz: p=5.6e^-78^). Similarly, at the population level, sensory stimulation led to a greater suppression of movement-responsive neurons during running but not resting **(Fig. 6c-d**, **Supplementary Fig. 5h**). Thus, sensory stimulation led to heterogenous changes of single neuron activities in a locomotion-state dependent manner. While the overall fraction of activated neurons was balanced by the fraction of inhibited neurons regardless of locomotion state, sensory stimulation robustly inhibited movement-related population dynamics during running.

**Figure 6:**
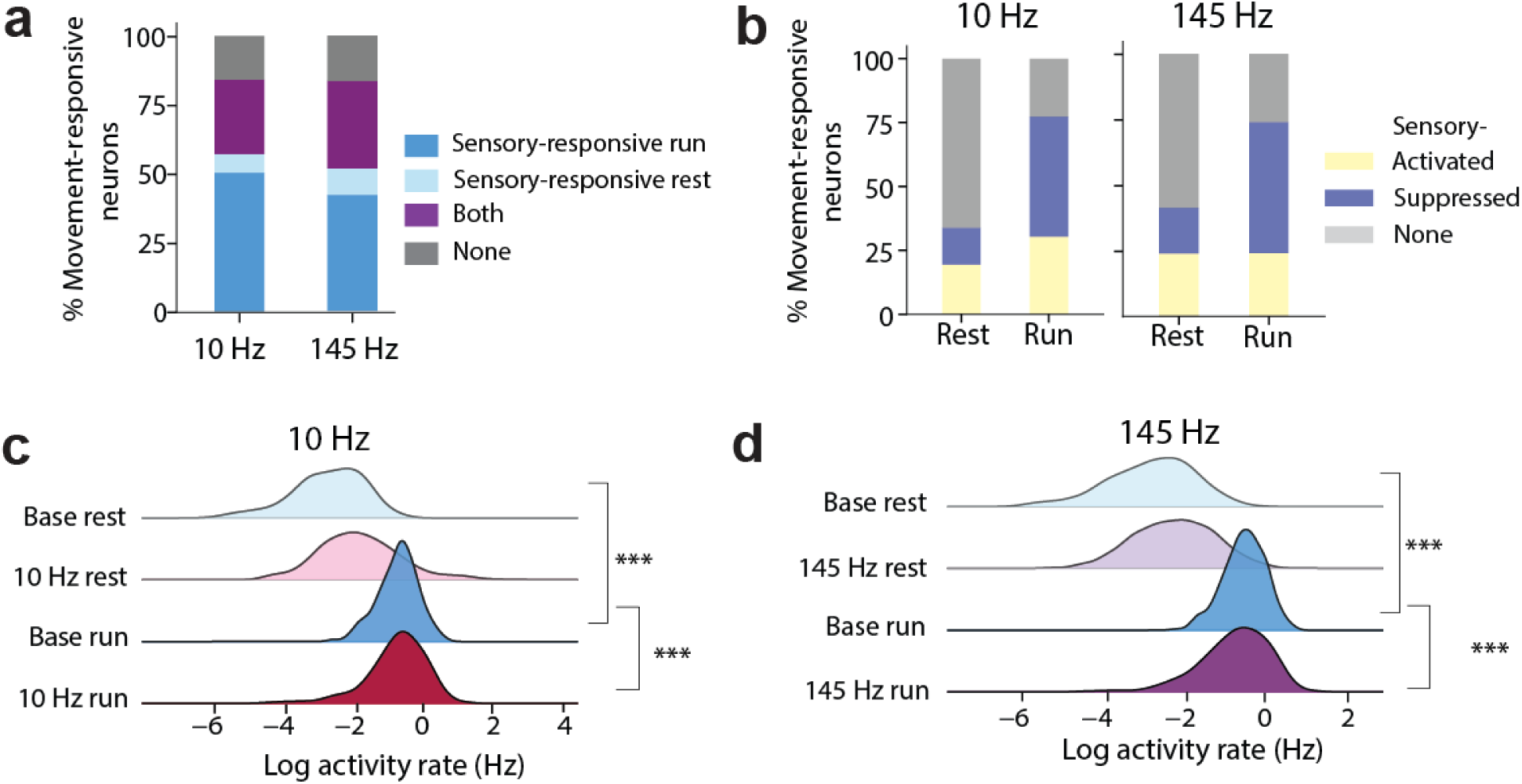
Sensory stimulation robustly suppressed movement-responsive neurons during running. **(a)** Stacked bar plot of the fraction of movement-responsive neurons that were responsive to 10 Hz (left) or 145 Hz (right) stimulation during resting (sensory resp. rest, light blue), during running (sensory resp. run, dark blue), during both resting and running (both, purple), and irresponsive (none, gray). **(b)** Stacked bar plot of the fractions of movement responsive neurons that were activated by 10 Hz (left) and 145 Hz (right) stimulation (activated, yellow), suppressed (suppressed, indigo) and non-modulated (none, gray), during resting vs. running. **(c)** Ridgeline plot of the population probability density of mean activity rate across movement-responsive neurons during resting or running, with vs. without 10 Hz stimulation (Mixed effect models, Baseline resting vs 10 Hz resting: p_approx_=1, Baseline running vs 10 Hz running: p_approx_=3.74e^-114^, n=2189). **(d)** same as (e) but during 145 Hz stimulation sessions (Baseline resting vs 145 Hz resting: p_approx_=1, Baseline running vs 145 Hz running: p_approx_= 4.6e^-158^, n=2041). *p < 0.05, **p < 0.01, ***p < 0.001.

### Both locomotion and sensory stimulation desynchronized movement-responsive neuronal networks

To further understand how sensory-evoked single neuron responses relate to network dynamics, we calculated pairwise Pearson correlations between movement-responsive neurons, as these neurons are most relevant to the locomotion behavior tested. We found that the overall correlation coefficients between these neurons across the entire recording sessions showed a negative linear relationship with the fraction of time the mice spent in running (**Fig. 7a**, R^2^=0.66, p=7.6e^-7^, n=25 sessions) and the average speed of the session (**Supplementary Fig. 6a)**, suggesting that locomotion generally desynchronizes the movement-responsive neural network, consistent with our previous study^45^.

**Figure 7:**
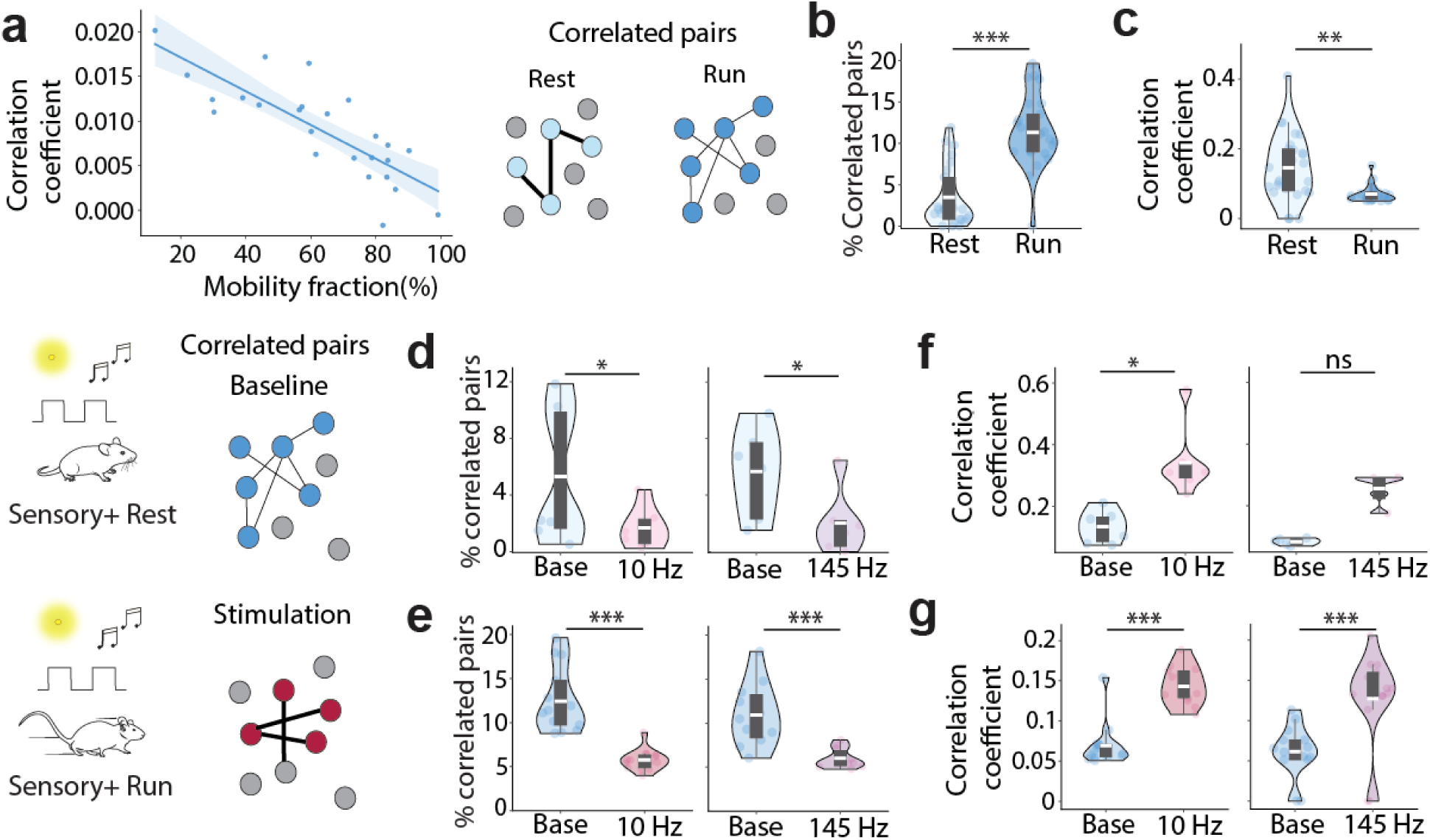
Locomotion and sensory stimulation both desynchronized the network of movement-responsive neurons. **(a)** Median correlation coefficients across movement-responsive neurons vs. the percentage of time mice spent running (mobility fraction). There was a significant linear relationship (linear regression, R^2^=0.66, p=7.6e^-7^, n=25 sessions) with shaded regions indicating the 95% confidence interval. **(b)** Percentage of correlated pairs during resting vs. running in the baseline period without stimulation (Wilcoxon signed rank test, p=2.3e^-5^, n=25). **(c)** Correlation strength of the correlated pairs during resting vs. running in the baseline (Wilcoxon signed rank test, p=0.002, n=25). **(d)** Percentage of correlated neuron pairs during baseline stimulation (Wilcoxon signed rank test, 10 Hz: p=0.016, n=7; 145 Hz: p=0.03, n=6). **(e)** Same as (d) but during running (Wilcoxon signed rank test, 10 Hz:1.2e^-4^, n=14; 145 Hz: 9.8e^-4^, n=11). **(f)** Quantification of correlation strength across correlated pairs during resting between baseline vs. stimulation (Wilcoxon signed rank test,10 Hz: p= 0.016, n=7; 145 Hz: p=0.06, n=6)**. (g)** Same as (f) but during running (10 Hz: p=2.4e^-4^, n=14; 145 Hz: p=4.9e^-4^, n=11). Quantifications are visualized as violin plots with the outer shape representing the data kernel density and a box plot (box: interquartile range, whiskers: 1.5x interquartile range, white line: mean). *p < 0.05, **p < 0.01, ***p < 0.001.

To further evaluate the impact of sensory stimulation and locomotion on functional connectivity, we identified neuron pairs that exhibited significant correlations given their corresponding activity rates. Briefly, we compared the correlation coefficient of a neuron pair to a shuffled distribution formed by assigning random lags between the neuron pair. If the observed correlation value was greater than 97.5^th^ percentile of the shuffled distribution (1000 shuffles), the neuron pair was deemed a ‘correlated pair’, otherwise a ‘random pair’. We found that the fraction of correlated pairs during running (run-relevant pairs) was significantly higher than during resting (rest-relevant pairs) (**Fig 7b**). Since the activity rates of these movement-responsive neurons increased during running (**Supplementary Fig. 6b)**, we computed the normalized correlation coefficient of each neuron pair by subtracting the median correlation of the corresponding shuffles. Interestingly, the normalized correlation coefficients among run-relevant pairs were significantly weaker than the rest-relevant pairs (**Fig. 7c**), which cannot be attributed to the activity rate differences between the populations (**Supplementary Fig. 6c-f**).

During sensory stimulation, the fraction of correlated pairs was significantly lower than baseline during both resting and running (**Fig. 7d, e**). Furthermore, stimulation enhanced the correlation strength of the correlated pairs compared to baseline, regardless of the movement state (**Fig. 7f, g**), which again cannot be explained by the variation in activity rates (**Supplementary Fig. 6c-f**). Together, these results demonstrate that both locomotion and sensory stimulation desynchronized the overall movement-responsive neural networks. However, locomotion was accompanied by an increased fraction of correlated pairs during movement, whereas sensory stimulation reduced the fraction, highlighting the distinct circuit effect of sensory versus motor processing in the striatum.

## Discussion

To understand the impact of beta-rhythmic sensory stimulation on stepping movement and basal ganglia circuits, we delivered audiovisual stimulation at either 10 Hz or 145 Hz to awake head-fixed mice voluntarily locomoting, while recording locomotion, dorsal striatal LFPs and cellular calcium dynamics. Consistent with the general ability for low frequency stimulation to entrain brain circuits, we found that sensory stimulation at 10 Hz effectively paced striatal population dynamics. While sensory stimulation at both frequencies promoted locomotion and desynchronized striatal network, only 10 Hz stimulation suppressed movement onset transitions, enhanced stepping rhythmicity at delta frequencies, and strengthened the coupling of LFP delta and beta power to the phase of delta-rhythmic stepping. Thus, beta-frequency sensory stimulation enhances gait rhythmicity by strengthening the interactions between striatal neural dynamics and stepping movement, distinct from the generally anti-kinetic effects observed with strong and artificial beta-frequency tACS or DBS electrical stimulation of cortical-basal ganglia circuits^30–35^. These results highlight the potential of beta-rhythmic sensory stimulation for improving gait via strengthening the coupling of striatal networks and stepping movement.

We found that 10 Hz stimulation resulted in a significant increase in 10 Hz LFP spectral power, which lagged the stimulation onset for ∼5.5 seconds and remained elevated for ∼8.5 seconds after offset (**Fig. 1g-h**). The long delay in LFP entrainment suggests a network mechanism that likely involves broad sensorimotor regions interconnected with the striatum^36,66^. Indeed, large-scale recordings revealed that visual stimuli evoked neural activity changes across many brain circuits in rodents^67,68^. Similarly, human EEG and fMRI studies demonstrated that sensory stimuli engaged broad brain regions, including the brainstem^69,70^. As LFPs are dominated by synaptic inputs, it is highly plausible that inputs from various cortical and subcortical areas contribute to the LFP entrainment effects observed in the striatum. While each brain region could support 10 Hz rhythms with distinct local circuit mechanisms, the coupling across cortical-basal ganglia-thalamic circuits most likely underlies the gradual strengthening of the 10 Hz striatal rhythm, resulting in the seconds’ long delay.

Stepping movement exhibits characteristic delta frequencies in both mice and humans^1,8^. Under our experimental conditions, mice movement rhythmicity peaked around 3-4 Hz. This delta component of movement speed was enhanced by 10 Hz sensory stimulation, but not 145 Hz stimulation (**Fig. 2c, e**), suggesting that 10 Hz stimulation leads to more rhythmic and consistent stepping patterns. These findings are supportive of the idea that 10 Hz sensory stimulation can engage delta-rhythmic neuronal activities via cross-frequency coupling that occur during natural physiological processes. This effect is distinct from the strong and artificial electrical or magnetic stimulation of the motor circuits that may evoke exaggerated oscillations which do not occur during physiological conditions^33–35^. Beta oscillations are conventionally thought to maintain status quo and once exaggerated, as in Parkinson’s disease, leads to akinesia^28^. We also observed that 10 Hz sensory stimulation lengthened running bouts (**Fig. 1p, Supplementary Fig. 2c**) and reduced movement onset transitions (**Fig. 1r**), consistent with the general notion that beta oscillations promote status quo and help maintain ongoing movement^25,27^.

Rodent whisking occurs around 8-16 Hz during active exploration^60,71,72^, which was found to be phase coupled to stepping in head-fixed mice^60^. We observed significant 10 Hz LFP entrainment during both running and resting, suggesting that stepping-coupled whisking had limited contribution to the LFP entrainment effect detected. Nonetheless, it is possible that 10 Hz sensory stimulation also entrains neural circuits related to whisking that further strengthens the observed LFP entrainment. Future analysis of the effect of sensory stimulation on whisking will help delineate whether sensory entrainment of rhythmic movement is a broad phenomenon, beyond the specific stepping movement examined here. We also observed prominent movement-dependent theta oscillations, which however showed little coupling to stepping rhythmicity (**Fig. 2f, g, i**), suggesting a potentially external origin, e.g. the hippocampus as previously demonstrated^73,74^.

We found delta-rhythmic stepping is synchronized with striatal LFP delta oscillations, which temporally organizes LFP beta and gamma power regardless of sensory stimulation (**Fig. 2i-k**). This observation is consistent with our previous findings^8^ and further confirms that dorsal striatum plays an important role in orchestrating stepping movement. Even though audiovisual stimulation at both 10 Hz and 145 Hz promoted movement, only 10 Hz stimulation strengthened the coupling of striatal LFP delta rhythms and stepping (**Fig. 2f, h**). This enhanced coupling likely arises from the entrainment of thalamic and cortical sensory inputs to the striatum. However, proprioceptive inputs due to delta rhythmic movement could also contribute to the strengthened coupling between LFP delta and stepping^2^.

It is intriguing that the faster beta oscillations augmented by sensory stimulation could modulate the slower delta oscillations to promote stepping automaticity. Neuronal delta rhythms and beta rhythms are temporally linked via cross-frequency coupling across motor circuits^2,28,75,76^, but generally with the slower delta rhythms setting up the temporal window to modulate the power of the faster beta rhythms. Our previous cellular voltage study showed that many striatal neurons exhibit delta rhythmic subthreshold membrane voltage fluctuations, with spiking occurring at beta frequency around the peak of membrane voltage delta oscillatins^8^. It is highly probable that the membrane biophysical mechanisms supporting the generation of delta and beta rhythms are linked, and thus selective enhancement of cellular beta rhythms could impact the slower delta rhythms. For example, beta rhythmic sensory inputs to a neuron could engage intrinsic ion channels involved in supporting cellular delta oscillations. Future voltage imaging studies that directly measure the cellular voltage effects of sensory stimulation could reveal insights on how higher beta frequencies impact the lower stepping-related delta frequencies at the individual neuron level.

We used GCaMP7f calcium imaging to probe the effect of sensory stimulation on individual striatal neurons and population striatal networks. To estimate neuronal spiking probability from cytosolic calcium that is directly measured by GCaMP7f, we identified calcium events as those with sharp rising phases and performed calcium event analysis using the onset or the entire rising phase. We were pleasantly surprised that calcium event onsets were phase-locked to the delta component of both stepping movement and LFP (**Fig. 4a-k**), despite the slow kinetics of calcium imaging, highlighting the ability of calcium imaging to track neuronal dynamics^8^. While striatal LFPs are dominated by presynaptic inputs, the observation that cellular calcium was phase coupled to LFP delta provides strong evidence that the striatal LFP delta rhythmicity organizes striatal outputs. The temporal relationship of calcium events with movement and LFP delta rhythms were largely unaffected by sensory stimulation, in agreement with the fact that calcium events reflect cellular spiking output probabilities, highlighting that sensory stimulation boosts locomotion without interfering with striatal locomotion encoding ability (**Fig. 4l-m**).

Interestingly, we found that ∼70% of neurons exhibited increase in cellular calcium during movement, similar to prior electrophysiological studies^61^, but greater than previous calcium imaging studies using GCaMP6f^43,45^, consistent with the improved sensitivity of GCaMP7f. Furthermore, consistent with previous electrophysiology studies, striatal cellular calcium rises preceded locomotion by approximately a hundred milliseconds^62^, supporting the role of striatum in regulating movement (**Fig. 3d-g**). We observed a sharp and transient increase of calcium activity at stimulation onset, and a transient rebound at stimulation offset (**Fig. 5a-i**). Upon sensory stimulation, individual striatal neurons exhibited diverse responses, with some increasing their activity rate and others decreasing. However, we found an equal proportion of activated and inhibited fractions across the population regardless of locomotion states (**Fig. 5k**), suggesting potential network regulation to maintain balanced excitability across behavioral states.

When focusing on movement-responsive neurons that are relevant for the locomotion task condition, we found that sensory stimulation has an inhibitory effect when mice were running, but not resting (**Fig. 6b-d**). It is possible that it is easier to detect inhibition when these movement-responsive neurons have higher activity rate during running in contrast to resting. Nonetheless, our results are consistent with previous studies showing sensory stimulation generates smaller responses in medium spiny projecting neurons (SPNs) during up-state than down-state in anesthetized mice^77^, and sensory suppression of spontaneous whisking-evoked striatal neuron activity in awake mice^62^. Previous studies also reported differential sensory responses between D1-SPNs and D2-SPNs^62,78^, though both SPN types tracked gait cycles^64^. As most of the neuron recorded here are SPNs, it will be interesting to examine whether some of the observed heterogeneity arises from the systemic difference between D1- versus D2-SPNs. In addition to SPNs, striatal cholinergic interneurons, known to be critical in regulating striatal dynamics may be robustly recruited by sensory stimulation. In particular, cholinergic drugs have been found to consistently improve gait stability in Parkinsonian patients, whereas dopamine replacement has mixed effects on gait and sometimes worsens gait^79,80^. Though it remains to be determined how much of the gait improvement effect of cholinergic drugs is mediated by striatal cholinergic interneurons versus a general enhancement of cognitive ability and sensorimotor integration^79,81^. Similarly, parvalbumin-positive fast spiking interneurons and other GABAergic interneurons have also been shown to be critical in sensorimotor behaviors^43^. Future cell type specific analysis will offer insights on how sensory stimulation engages different striatal neuron subtypes and their involvement in gait regulation.

At the network level, we found that sensory stimulation and locomotion both desynchronized movement-responsive neuron populations, though through distinct effects (**Fig. 7**). During locomotion, we observed an increase in the proportion of correlated neuron pairs, similar to previous findings of increased functional connectivity within neuron clusters during movement^82^ (**Fig. 7b**). However, the connectivity strength between these correlated neuron pairs was lower during running compared to resting, pointing to a desynchronization effect (**Fig. 7c**). Desynchronization increases network information coding, which would be consistent with elevated processing of the environment as animals move^83^, even though under our experimental condition mice were head fixed. During stepping, although striatal neurons’ calcium activity rates increased, we detected a reduction in pairwise correlation coefficients that may stem from greater asynchrony between calcium event timing across neurons. Such asynchrony could arise from many mechanisms, including increased membrane conductance of neurons that render them less sensitive to synaptic inputs, recruitment of inhibitory cells, non-overlapping or desynchronized inputs to striatal neurons, lateral inhibition between striatal neurons, or temporally jittered spiking outputs to membrane depolarizations^84–86^. During sensory stimulation, the heightened sensory inputs to striatum could engage similar cellular and network mechanisms as during locomotion to enhance network information coding by reducing neuronal synchrony. While sensory stimulation reduced the fraction of correlated cell pairs in the movement-responsive striatal network, the correlation strength within these neurons increased (**Fig. 7d-g**), suggesting that sensory inputs have the potential to synchronize specific neuron populations receiving joint inputs. Overall, the sensory stimulation evoked desynchronization supports a potential therapeutic benefit in boosting movement by making the striatal network more dynamic and flexible.

There is a natural decline in gait automaticity with aging, and in particular Parkinson’s disease is characterized by various gait impairments, including heightened temporal variability and asymmetry, diminished rhythmicity, and shortened step length^19,80^. The malfunction of basal ganglia circuits in Parkinson’s disease is thought to disrupt habitual or automatic process^87^, resulting in heightened cognitive effort during stepping and loss of gait automaticity. To compensate for the diminished gait automaticity, Parkinsonian patients often rely on sensory cues during walking^14–16^, as sensory cue may help regularize input rhythmicity to provide a temporal framework during stepping^88,89^. Further, it has been suggested that abnormal neuronal synchrony in Parkinson’s disease may result from a loss of active decorrelation within the basal ganglia^45,90^, and our previous study linked abnormal increase in striatal synchrony to motor deficits in low- dopamine conditions^45^. The observation that sensory stimulation desynchronizes striatal network also supports a desynchronization mechanism underlying heightened locomotion during sensory stimulation. Given the non-invasive nature of sensory stimulation, our study highlights an exciting translational potential of using sensory stimulation to influence various aspects of movement, including gait rhythmicity, the propensity to move, and movement onset transitions particularly in patients with movement and gait disorders.

## Methods

### Animal Surgery

All animal procedures were approved by the Boston University Institutional Animal Care and Use Committee. The study included both female and male C57BL/6 mice (n=9, 7 females and 2 males) 8-12 weeks at the start of the experiments, obtained from Charles River Laboratories. Data from male and female mice were pooled. Mice were first surgically implanted with a sterilized custom imaging window over the dorsal striatum (anteroposterior 0.5mm, mediolateral 1.8mm, dorsoventral -1.6mm from the brain surface), a small ground pin over the cerebellum as the LFP recording reference, and a headplate. The imaging window consisted of a stainless-steel cannula (outer diameter 3.17mm, inner diameter 2.36mm, height 2mm) adhered to a circular coverslip (size 0, outer diameter 3mm) with ultraviolet curable optical adhesive (Norland Products), coupled to an infusion cannula (26 gauge; No.C135G-4; Plastics One) and an insulated stainless-steel wire (Diameter: 130 µm, Plastics One Inc., 005SW-30S, 7N003736501F) for LFP recordings. The stainless-steel wire was glued to the infusion cannula and protruded from the bottom of the infusion cannula by about 200μm.

To image the dorsal striatum, the overlying cortical striatal tissue was gently aspirated to gain optical access^43,45,91^. While the tissue removal was necessary and not ideal, this animal preparation did not result in any noticeable behavioral deficits, including rotational behaviors as reported previously^45^. Following recovery from surgery (14-21 days post-surgery), one cohort of mice (n=4) were injected with a 1μl AAV9-Syn-GCaMP7f.WPRE.SV40 virus (titer 6.6 x 1012 GC/mL) through the attached guide cannula. The other cohort (n=5) was injected with a 1μl AAV9-Syn- SomaGCaMP7f.WPRE.SV40 virus (titer 6.6 x 1012 GC/mL). After viral infusion (21-28 days), animals were habituated to voluntary locomotion on a spherical treadmill for 2-3 days per week for 2 weeks. Each habituation session lasted for ∼30 min per day, in a dimly lit room under similar conditions as during recording sessions.

### Locomotion, imaging and LFP data acquisition

During each recording session, mice were awake, head fixed above a spherical treadmill that permitted voluntary locomotion. The spherical treadmill consisted of a Styrofoam ball supported by pressurized air (∼50 psi) in a custom 3D printed plastic housing as in our previous studies^43,45,92^. Movement was monitored with two computer universal serial bus sensors fixed to the plastic housing at an angle of 75 degrees at the equator. The x- and y- displacement acquired by each sensor was collected at 20 Hz using a custom Teensy system and recorded via a custom MATLAB script as described in Romano et al^92^.

Image acquisition was carried out using a custom microscope equipped with a scientific complementary metal oxide semiconductor (sCMOS) camera (ORCA-Flash4.0 LT Digital CMOS camera, No. C11440-42U, Hamamatsu), a Leica N Plan 10x0.25 PH1 microscope objective lens, an excitation filter (No. FF01-468/553-25), a dichroic mirror (No. FF493/574-Di01-25x36), an emission filter (No. FF01-512/630-25; Semrock) and a tube lens (Nikon Zoom-NIKKOR 80-200 mm f/4 AI-s). GCaMP7f fluorescence was excited by a 5 W light-emitting diode (<u>LZ1-00B200,</u> <u>460 nm; LedEngin</u>). The imaging field of view was 1.343 x 1.343 mm^2^, with each pixel corresponding to 1.312 x 1.312 μm^2^. Image acquisition was though HC Image Live (Hamamatsu) at 20 Hz. Image data were stored in dcimg format and converted into tiff format for offline processing.

LFPs were recorded with OmniPlex (PLEXON) at 1 kHz. To synchronize movement data, calcium imaging data, LFPs and sensory stimulation, OmniPlex also recorded the TTL pulses used for each sCMOS image frame acquisition, Teensy motion sensor reading, and audiovisual sensory stimulation triggers.

### Sensory stimulation

Visual stimulation was delivered by a white LED placed ∼5cm away from the eye. Auditory stimulation was through a speaker within earshot at ∼77 decibel. Both light and sound pulses were delivered at 50 % duty cycle via a function generator (33220A, Keysight). Each recording session consisted of five trials of stimulation at either 10 Hz or 145 Hz. Only one stimulation frequency was used per day. Each trial contains a one-minute-long sensory stimulation, followed by five minutes of baseline without stimulation (**Fig. 1b**). There was also 6 minutes of baseline period before the start of the first trial, resulting in a total of 36 minutes per recording session. One or two sessions were recorded for each mouse for each stimulation frequency.

### Movement speed trace extraction

Animal’s movement speed was computed as the linear velocity of the treadmill movement using the following formula:

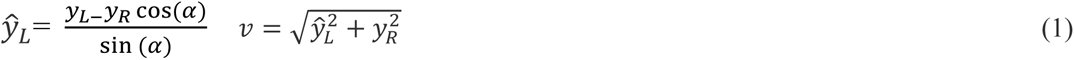

where α is the angle between the two sensors (75 degrees), v is the instantaneous velocity of the mouse, and y_L_, y_R_ are the vertical readings from the left and right sensors as in Romano et al^92^. We then identified sensor artifacts as points greater than 100 cm/s and removed these artifacts by replacing them using linear interpolation.

***Identification of rest period and running bouts:***

We first down-sampled the movement speed trace using a moving average with a window size of 20 frames (1 second) (movmean in MATLAB). Since the variance of speed during resting is low, we computed the variance of the down-sampled speed trace using a 2 second window (40 frames) and identified “putative stationary periods” as periods with variance less than 0.1, a mean speed less than 5 cm/s and a duration longer than 2 seconds. We then computed the low-speed threshold as the mean + 2 standard deviations of the speed during the putative stationary periods. For two sessions in which mice had extensive movement, the putative rest periods could not be reliably estimated, and thus we set the low-speed threshold to 2.5 cm/s based on manual inspection of the speed traces.

Using the low-speed threshold, we then identified rest periods utilizing a sigmoid function. Specifically, we first assigned each movement speed data point (v) a value using the function F (equation 2), which scales all points higher than the threshold towards zero.

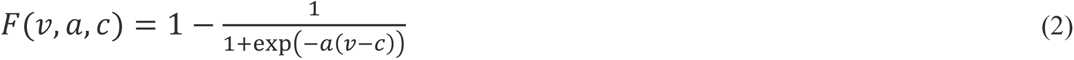

where v is the movement speed trace, c is the low-speed threshold, a is set to 0.8 (sigmoid slope). We then computed the scaled speed trace F using a moving window of 2 seconds. Data points of the scaled speed trace F greater than 0.45 for at least 2 seconds were deemed to be the rest periods. Note, each rest period was at least 2 seconds long.

Similarly, running bouts were identified using the sigmoid function defined in equation 3. The scaled speed trace G was computed using a moving window of 2 seconds. Data points of the scaled speed trace G greater than 0.55 for at least 2 seconds were deemed to be the running bouts. Note, each running bout was at least 2 seconds long.

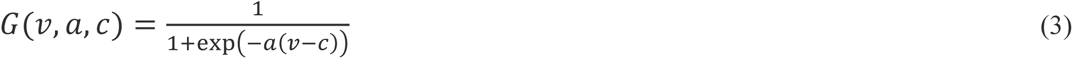

### Movement onset transition identification

Movement onset transition was defined as the onset of running bouts preceded by resting periods. To determine the onset point of each running bout, we first computed a high-speed threshold for each session as the 50^th^ percentile of the speed during all periods other than resting periods. A “putative onset point” is then defined as the first time point within each running bout where movement speed exceeded the value of the low-speed threshold + 10% of the difference between the high-speed threshold and the low-speed threshold. We then determined the exact onset point using a search-box to find the time point within 2 seconds before and after the putative onset point, where the acceleration exceeded threshold (high speed threshold-low speed threshold)/3) and stayed above the threshold for at least 250 ms. Visual inspection was performed to confirm the accuracy of the onset detection.

### Movement speed rhythmicity analysis

Movement speed rhythmicity was only examined during running bouts. Stepping rhythmicity was estimated by calculating the delta frequency power of the movement speed trace. Specifically, movement speed trace was first band pass filtered using a Butterworth filter in the delta frequency range (3-4 Hz), and then Morelet wavelets power spectrograms were calculated with the FieldTrip toolbox^93^ (https://www.fieldtriptoolbox.org/) for Matlab.

### Calcium imaging pre-processing, ROI segmentation and calcium event detection

Each calcium imaging video was first motion corrected as detailed in Tseng et al^94^. We then calculated a maximum fluorescence minus minimum fluorescence projection image of the motion- correct image stack, and manually identified regions of interests (ROIs) corresponding to individual cells as circles with a radius of 6 pixels (7.8 µm). The ROI intensity trace was computed as the mean intensity across all pixels within each ROI. A local background ‘donut’ trace was calculated as mean intensity of all pixels within 50 pixels of the centroid of the ROI excluding pixels within any identified ROIs. This resulting background trace was further subtracted from the corresponding ROI’s trace to minimize tissue fluorescence scattering. Next, the fluorescence traces (Δf/f) for each neuron were computed as ROI intensity trace minus the local background ‘donut” trace, normalized by the mean across the entire session. GCaMP7f fluorescence traces were then interpolated to exactly 20 Hz and linearly detrended for every neuron. All traces were visually inspected before further analysis. The last minute (35-36 min) of all recording session was excluded for subsequent analyses as some sessions showed evident photobleaching.

Calcium event detection was based on Friedrich et al^95^, which uses a generalized pool adjacent violators algorithm (PAVA) to infer the most likely event train. Specifically, we used a threshold that is 2.5 times of the standard deviation of the detrended fluorescence trace on the discretized deconvolved output. We performed a manual inspection of detected events in about 100 traces from multiple mice to ensure the accuracy of event detection. To generate calcium activity traces, the rising phase of each calcium event was binarized as ones, and the rest as zeros. Calcium activity rate across all recorded neurons was 18.39 ± 3.69 frames/min at 20 Hz frame rate (mean ± std dev, n = 6096 neurons from 9 mice).

### Identification of movement responsive neurons

A neuron was categorized as movement responsive if its calcium activity rate was significantly higher during running than resting. Specifically, we concatenated all the rest or run periods in baseline and computed the difference between the calcium activity rate during running versus resting (D).

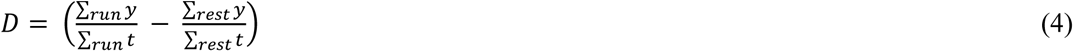

A shuffled distribution was created by randomly selecting equivalent running and resting periods from the entire baseline period 1000 times and calculating the difference in each shuffle. If the observed difference was greater than 97.5^th^ percentile of the shuffled distribution, the neuron was classified as a movement responsive neuron. Two sessions that had less than a minute of rest periods in baseline were excluded from this analysis due to the lack of sufficient baseline for estimating the shuffled distribution.

### Cross correlation analysis between population activity rate and movement

We estimated the correlation coefficient between the speed trace and the summed calcium activity rate across movement responsive neurons at different time lags using the python package scipy.correlate. Significance in correlation was compared to the shuffled distribution built by temporal shuffling (1000 times) the summed calcium activity rate relative to the speed trace by random lags and calculating the corresponding correlation coefficient. The temporal lag was estimated as the timepoint corresponding to the peak correlation coefficient. Sessions with lags longer than 1 second were excluded from this analysis.

### Detection of stimulation responsive cells

We identified stimulation responsive neurons by comparing the mean activity rate during stimulation versus baseline in the same locomotion state (rest or run). The shuffled distribution (1000 shuffles) was built by computing the mean activity rate during randomly chosen baseline period in a particular locomotion state during each shuffle. If the observed value during stimulation period was significantly higher than the 97.5^th^ percentile of the shuffled distribution, the neuron was considered activated by sensory stimulation. Similarly, if the activity rate was lower than 2.5^th^ percentile of the shuffled distribution, the neuron was deemed inhibited. Sessions with less than one minute of stimulation period in a given locomotion state or if the duration of baseline period was less than twice the duration of the corresponding stimulation period were excluded for this analysis.

### Neuron pairwise correlation Analysis

Correlation coefficient was calculated as the Pearson correlation coefficient of a neuron pair’s calcium activity traces. A shuffling procedure was used to determine whether a neuron pair was significantly correlated. For each shuffle, we shifted the calcium activity traces of the neuron pair by a random delay and computed the corresponding correlation coefficients. This process was repeated 1000 times to build a shuffle distribution for every neuron pair. If the observed value was greater than 97.5^th^ percentile of the distribution, the neuron pairs were deemed ‘correlated pairs’. Due to the sparsity of calcium events, negatively correlated pairs (<2.5^th^ percentile) were excluded for subsequent analyses.

To evaluate the modulation of correlation during specific periods of interest, we concatenated activity traces in those periods and computed the median correlation coefficient across the correlated neuron pairs. To estimate the impact of event rate on the identification of correlated pairs, we grouped neuron pairs by onset frequencies and computed the threshold (97.5^th^ percentile) of the shuffled distribution and found the threshold had no correlation with activity rate (**Supplementary Fig. 7b**). Similarly, to estimate the impact of activity rate on correlation coefficients, we examined the mean and median correlation coefficient for neuron pairs with different onset frequencies (**Supplementary Fig. 7a**, c, d). The median across shuffles increases with high activity rates (**Supplementary Fig. 7c**). Thus, we computed the correlation coefficients by subtracting the median correlation coefficient across 1000 shuffles from the observed correlation coefficient for every neuron pair to correct for any bias due to activity rate difference.

### LFP power spectrum

Intrinsic to awake recordings, LFP recordings were occasionally affected by motion artifacts. We first excluded potential motion artifacts, points greater than mean ± 5 standard deviations. The excluded periods were less than 1 minute for each of the 36 minutes long recording session, except one recording session that was completely excluded due to long-lasting artifacts. LFP power spectrum was computed using Complex Morlet wavelet transformation with Pywavelets package^96^ (‘cmor14.0-1.5’ with bandwidth 14, center frequency 1.5, and scaled between 5 and 5000). The power spectrogram was further normalized by the total power across all frequencies at each time point to probe the relative distribution of power at a specific frequency band. LFP power for theta (6-8 Hz), beta (15-30 Hz) and gamma (45–80 Hz) bands were computed by averaging the power across all frequencies within these bands. To compare the power between running and resting periods during baseline without stimulation, we combined baseline periods of 10 Hz and 145 Hz recording sessions.

To determine LFP power change around movement delta rhythmicity (**Fig. 2l**), we first aligned LFP power spectrogram to each movement delta oscillation peak and normalized LFP power to the mean power across the given delta-peak triggered window. We then averaged the normalized LFP power spectrogram across all delta peaks in all recording sessions to generate the movement speed delta oscillation peak triggered LFP spectrogram.

### Phase locking analysis

We first applied Hilbert transform to find the instantaneous power at each frequency. The speaker system produced artifacts in some sessions that can be easily identified as a sharp and consistent change in LFP waveforms during each stimulation. We excluded these sessions from phase locking analysis by manually inspecting each individual LFP trace, the average LFP waveform triggered by all stimulation pulses of a session, and the LFP power spectrum.

To compute phase locking value (PLV) between LFP and stimulation pulses, we used the following equation:

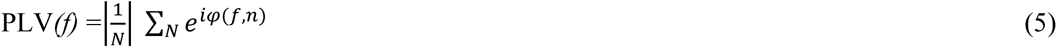

where the phase of a given frequency *f* was obtained from the Hilbert Transform of the stimulation pulse train using a Butterworth filter (filter order = 2) around 10 Hz.

To compute the PLV between LFP and movement, we used the same equation (5) where we bandpass filtered movement speed traces and LFP traces at delta frequencies (2–4 Hz) using a Butterworth filter and calculated the instantaneous phase using Hilbert Transform.

To compute the phase-locking of calcium event onsets to movement speed (**Fig. 4a-g, 4l**) and LFPs (**Fig. 4a-c, h-k, m**), we first bandpass filtered movement speed traces or LFP traces at delta frequencies (2–4 Hz) using a Butterworth filter and calculated the instantaneous phase using Hilbert Transform. Since PLV is biased by low event numbers, we utilized an adjusted phase locking value (PLVu^2^) that accounts for the number of events and only included neurons with at least 20 events. PLVu^2^ were computed using the following equation as in our previous study^8^:

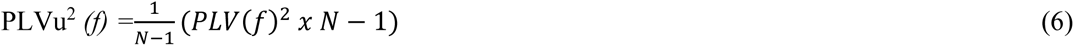

where N is the number of calcium events. To test for statistical significance, a shuffled distribution for each neuron was generated by computing PLVu^2^ with randomized event times. We then used mixed effect model between the observed value and the shuffled distribution.

Phase angle polar histograms were created by plotting the preferred phase (circular mean value) of each neuron relative to the delta-filtered movement speed or LFP signals.

### Cross-frequency coupling (CFC) between movement speed and LFP

We first computed the instantaneous phase of movement speed delta oscillation using Hilbert transform from the delta filtered movement speed trace (Butterworth filter, filtered order=2). Similarly, we computed the instantaneous amplitude of LFPs across beta/gamma frequencies, and then derived the instantaneous phase of beta or gamma amplitude fluctuations. CFC of movement speed delta and LFP beta/gamma were computed as the PLV between these phase vectors.

### Statistical analyses

All comparisons were made with versus without sensory stimulation in the same mice to rule out any potential impact due to variation of animal preparation procedure between mice. To compare LFP and movement changes between stimulation and baseline, we used Wilcoxon signed rank test, with a significance level α = 0.05 across trials or sessions. To compare fractions, we used Fisher’s exact test with error bars computed as the 95% confidence interval defined as^83^:

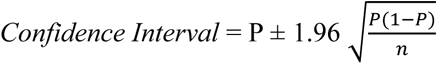

where *P* denotes the percentages, and *n* denotes the number of samples.

To compare changes involving individual neurons, we used Mixed effect models (MATLAB function fitglme, Poisson distribution with log link function). The fixed effects in the model were movement speed (rest and run), sensory stimulation (on and off) and an interaction term (speed x stimulation). The random effect was session IDs and neurons.

To determine neuronal calcium activity rate changes during stimulation, we used the following model:

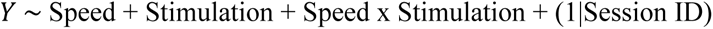

where Y is the calcium activity rate. We computed the R^2^ for the model and tested the coefficients for significance (10 Hz: R^2^ =0.59, n=2189; 145 Hz: R^2^=0.49, n=2041). We also compared the fit of the model to an intercept-only model, speed-only model, speed, and stimulation without interaction model using log likelihood and Akaike information criterion (AIC). Higher AIC with lower log likelihood indicates better fit of the model. We confirmed that the chosen model exhibited highest AIC and lowest log likelihood compared to other models tested. Using this model, speed was identified as a significant factor in the absence of sensory stimulation (mixed effect model, 10 Hz: p_approx_=0, 145 Hz: p_approx_=0), confirming that movement responsive neurons showed a robust increase in activity rate during sustained locomotion.

To test for the influence of stimulation on PLVu^2^ between baseline and stimulation during running, we used the following mixed effect model:

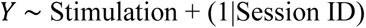

where Y is the PLVu^2^. As the preferred phase is sensitive to both dispersion and direction, we assessed statistical significance using the Kuiper’s test (circ_cmtest), which is a circular analogue of the Kruskal-Wallis test.

## Author contributions

S.S. and X.H. designed the study. S.S conducted the experiments, collected and analyzed the data. E.L. assisted with data analysis. H.J.G. and J.F performed mouse surgeries and assisted with data collection. F.L. helped with data collection. C.Z assisted with data analysis. S.S. and X.H. prepared the manuscript, and all authors edited the manuscript. X.H. oversaw all aspects of the project and supervised the study.

## Data Availability

Small datasets to demo the software/code are available at the Github repository https://github.com/HanLabBU/Sridhar-Sensory-Modulation-Striatum (Note, final link will be updated upon publication). Additional experimental data are available from the lead contact upon request.

## Code Availability

Code used for data analysis is currently available at the Github repository https://github.com/HanLabBU/Sridhar-Sensory-Modulation-Striatum

## Acknowledgements

We thank members of Han Lab for providing invaluable help throughout the study, and Dr. Mark Kramer for guidance on statistical tests. X.H. acknowledges funding NIH 1R01NS115797, NSF CBET-1848029 and NSF CIF-1955981.

