## Supplemental Info for "Beta-frequency sensory stimulation enhances gait rhythmicity through strengthened coupling between striatal networks and stepping movement"

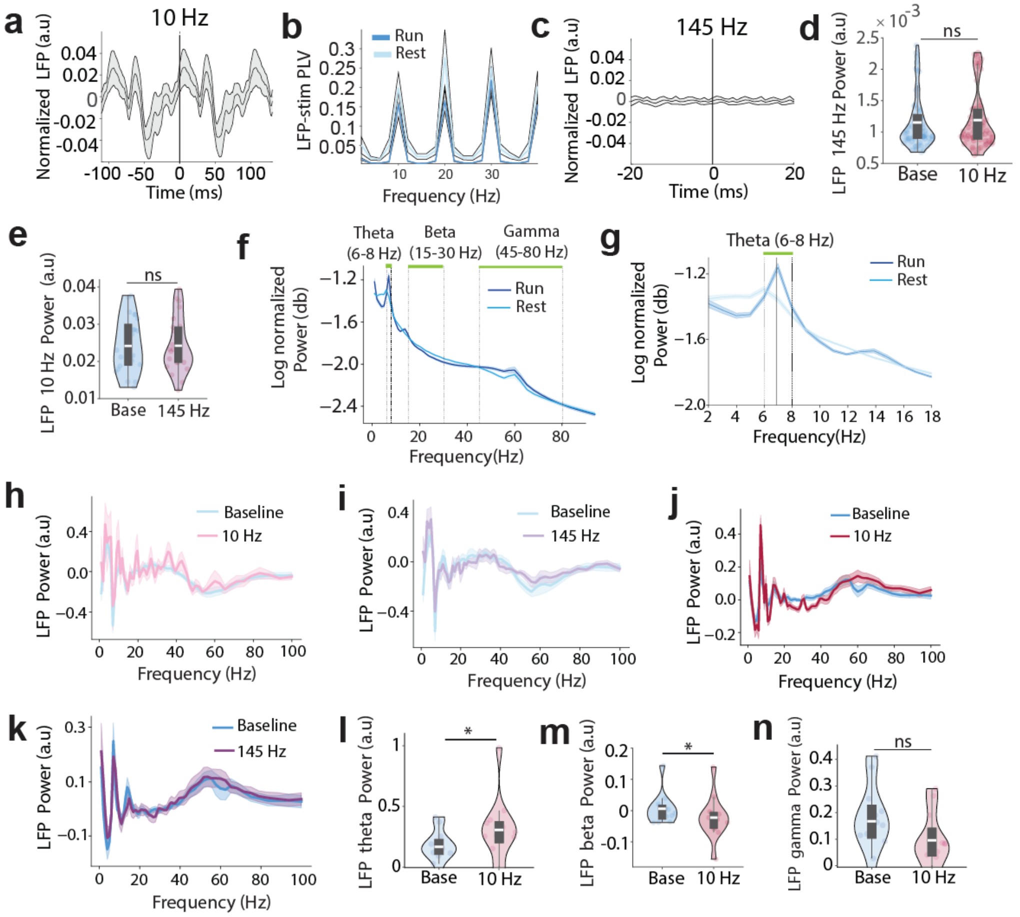


**Supplementary Figure 1: (a)** Population LFPs aligned to 10 Hz stimulation onset. Line: mean across trials; shaded region: standard error of the mean. **(b)** Population phase-locking value (PLV) between 10 Hz stimulation pulse trains and LFP across frequencies during running (dark blue) and resting (light blue). Line: mean across trials; shaded region: standard error of the mean. There was no difference in the PLV at 10 Hz between the resting and running conditions (Wilcoxon signed rank test, 10 Hz: $p=0.4$, $n=65$ trials). **(c)** Same as (a) but aligned to 145 Hz stimulation onset. **(d)** LFP 145 Hz power during the 1-minute period pre- (base) versus post- 10 Hz stimulation (Wilcoxon signed rank test, $p=0.12$, $n=65$ trials). **(e)** LFP 10 Hz power during the 1-minute period pre- (base) versus post- 145 Hz stimulation onset (Wilcoxon signed rank test, $p=0.75$, $n=24$ trials). **(f)** Normalized LFP power spectrum density (PSD) during running (dark blue) and resting (light blue) in baseline periods across all trials. Shaded region indicates standard error of the mean. **(g)** A zoom-in of (**f**). **(h)** Normalized LFP PSD during resting in baseline (blue) and 10 Hz stimulation (red) **(i)** Same as (**h**) but for 145 Hz stimulation (purple). **(j)** Same as (**h**), but during running in baseline versus 10 Hz stimulation. **(k)** Same as (**j**) but for 145 Hz stimulation. **(l)** Change in LFP theta power during 10 Hz stimulation compared to baseline when mice were running (Wilcoxon signed rank test, $p=0.034$). **(m)** Change in LFP beta power during 10 Hz stimulation compared to baseline when mice were running (Wilcoxon signed rank test, $p=0.034$). **(n)** Change in LFP gamma power during 10 Hz stimulation compared to baseline when mice were running (Wilcoxon signed rank test, $p=0.5$). Quantifications are visualized as violin plots with the outer shape representing the data kernel density and a box plot (box: interquartile range, whiskers: 1.5x interquartile range, white line: mean). $*p < 0.05$, $**p < 0.01$, $***p < 0.001$.

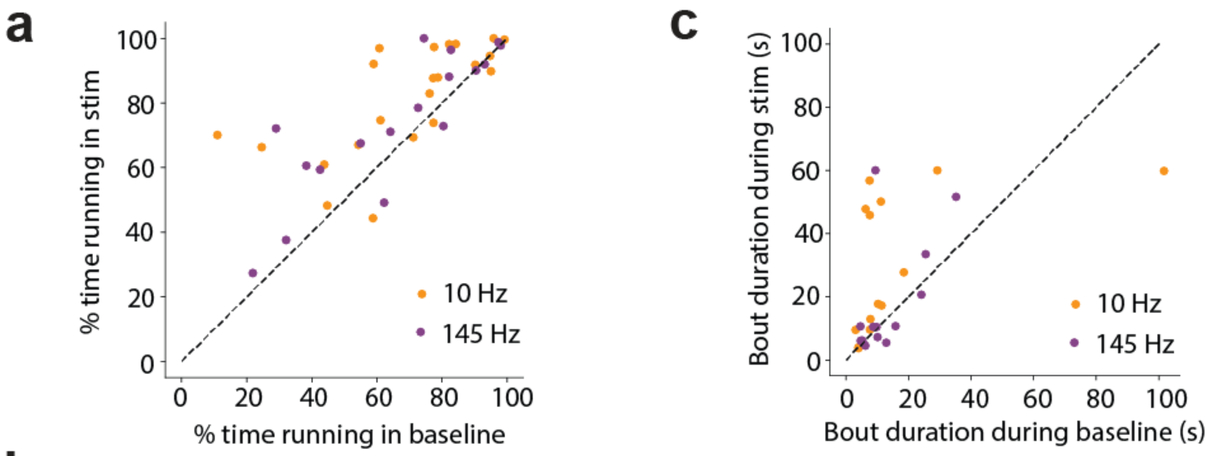


**b**

| Frequency | Condition | % Running period | Speed (cm/s) |
|---|---|---|---|
| 10 Hz | Baseline | 69.6 ± 22.3% | 15.7 ± 10.3 (0.5 - 58.6) |
| | Stimulation | 82.2 ± 16.5% | 14.6 ± 8.5 (0.5 - 55) |
| 145 Hz | Baseline | 65.7 ± 24.4% | 16.1 ± 8.8 (0.6 - 49.7) |
| | Stimulation | 74.1 ± 21.3% | 13.7 ± 7.9 (0.7 – 44.2) |

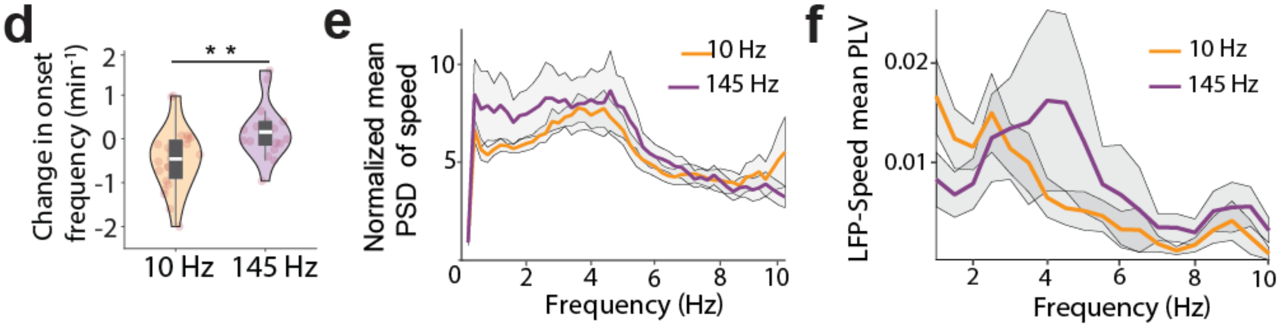


**Supplementary Figure 2: (a)** The percentage of time mice spent running during 10 Hz (orange) and 145 Hz stimulation (purple) vs. during baseline without stimulation. Each dot corresponds to a session. There was no difference between 10 Hz and 145 Hz during baseline (Wilcoxon rank sum test, p= 0.61) or stimulation (Wilcoxon rank sum test, p= 0.2). **(b)** Table indicating the percentage of time mice spent running and the corresponding speed during various experimental conditions. Mice ran 11-98% of the time across recording sessions, with a mean of 67.9 ± 23.3%, and a speed of 15.7 ± 10.3 cm/s (0.5 **-** 58.6 cm/s). **(c)** Running bout durations during 10 Hz (orange) and 145 Hz stimulation (purple) vs during baseline. Each dot corresponds to the median of a session. Bout duration during 10Hz stimulation was significantly longer than during baseline (Wilcoxon signed-rank test, p=0.02). There was no change during 145 Hz stimulation relative to baseline (Wilcoxon signed-rank test, p=0.5). **(d)** Stimulation evoked changes in movement onset transition frequencies (stimulation-baseline) at 10 Hz and 145 Hz (Mann Whitney U test, p=0.002, 10 Hz: n=23 sessions, 145 Hz: n=17 sessions). **(e)** Mean PSD of movement speed during running across all baseline sessions tested for 10 Hz versus 145 Hz stimulation. There was no difference (Wilcoxon Rank-sum test, p=0.84). **(f)** Mean LFP-movement speed PLV during running across all baseline sessions tested for 10 Hz versus 145 Hz stimulation. There was no difference (Wilcoxon Rank-sum test, p=0.96).

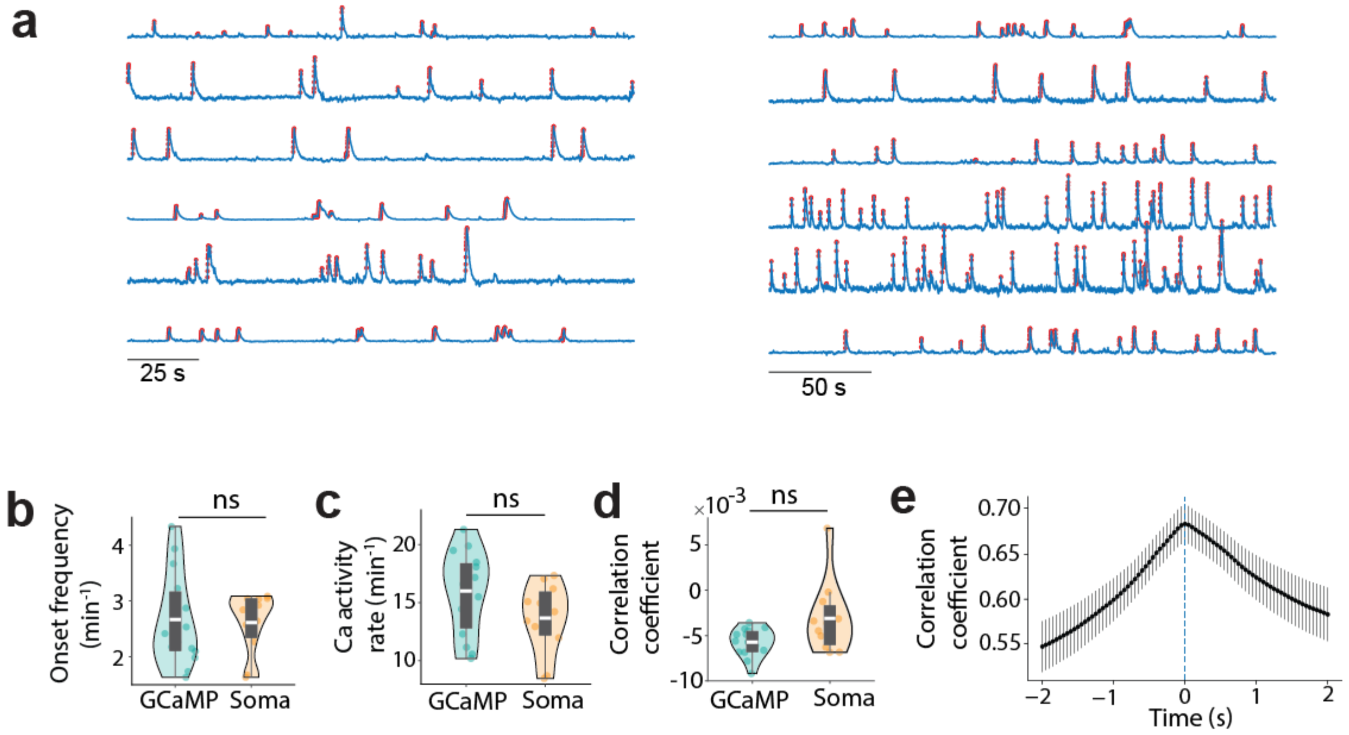


**Supplementary Figure 3: (a)** Example GCaMP7f traces from representative neurons during a typical recording session. The rising phases of identified events were marked with red. **(b)** Mean calcium event onset frequencies across all GCaMP7f baseline recording sessions without any stimulation vs SomaGCaMP7f baseline recording sessions. Each dot corresponds to the mean event frequencies across all neurons in a session. There was no difference (GCaMP7f: 2.67 ± 0.79 events/min, n=15 sessions, somaGcamp7: 2.61 ± 0.51 events/min, n=11, Independent t-test, p=0.86). **(c)** Mean calcium event activity rate across all GCaMP7f baseline recording sessions versus SomaGcaMP7f baseline recording sessions. There was no difference (GCaMP7f: 16 ± 3.41 /min, n=15, SomaGcaMP7f: 13.67 ± 2.87/min, n=11, Independent t-test, p=0.08). **(d)** Median correlation coefficients across all GCaMP7f baseline recording sessions versus SomaGCaMP7f baseline recording sessions (Wilcoxon rank sum test, p=0.052, n=15 GCaMP7f sessions and n=11 SomaGCaMP7f sessions). **(e)** Population correlation coefficient across time lags, aligned to peak correlation coefficient across all recording sessions. Quantifications are visualized as violin plots with the outer shape representing the data kernel density and a box plot (box: interquartile range, whiskers: 1.5x interquartile range, white line: mean). *p < 0.05, **p < 0.01, ***p < 0.001

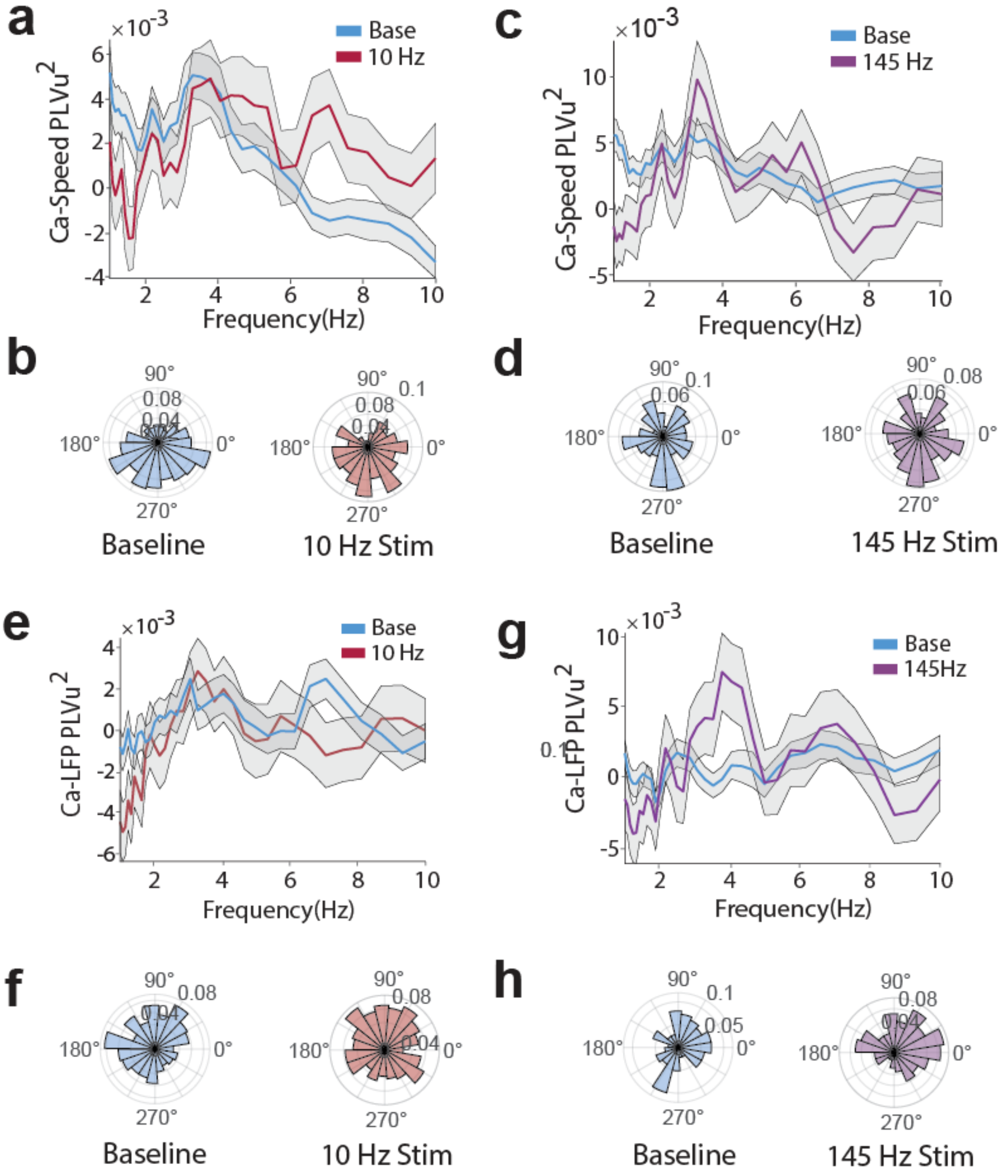


**Supplementary Figure 4: (a)** Population PLVu$^2$ of calcium event onsets to movement speed across frequencies during 10 Hz stimulation (red) and baseline (blue). There was no difference in delta frequency (2-4 Hz) phase locking strength (Mixed effect models, p= 0.27, n=907 neurons). **(b)** The polar histogram of neurons' preferred calcium event phase (mean angle) to delta-filtered speed during 10 Hz stimulation (red) and baseline (blue). There is no difference (Kuiper's Test, p=0.8). **(c, d)** Same as (a, b) but for 145 Hz stimulation (purple) (PLVu$^2$, mixed effect models, p= 0.25, n=659 neurons; preferred phase, Kuiper's test, p=0.7). **(e)** PLVu$^2$ of calcium event onsets to LFP across frequencies during 10 Hz stimulation (red) and baseline (blue) across all neurons. There was no difference in delta frequency (2-4 Hz) PLVu$^2$ (Mixed effect models, p= 0.44, n=889 neurons). **(f)** The polar histogram of the phase of calcium event onsets to LFP delta oscillations during 10 Hz stimulation (red) and baseline (blue). There is no difference (Kuiper's Test, p=0.2). **(g, h)** Same as (e, f) but for 145 Hz stimulation (purple) (PLVu$^2$, mixed effect models, p=0.32, n=676 neurons; preferred phase, Kuiper's Test, p=0.1).

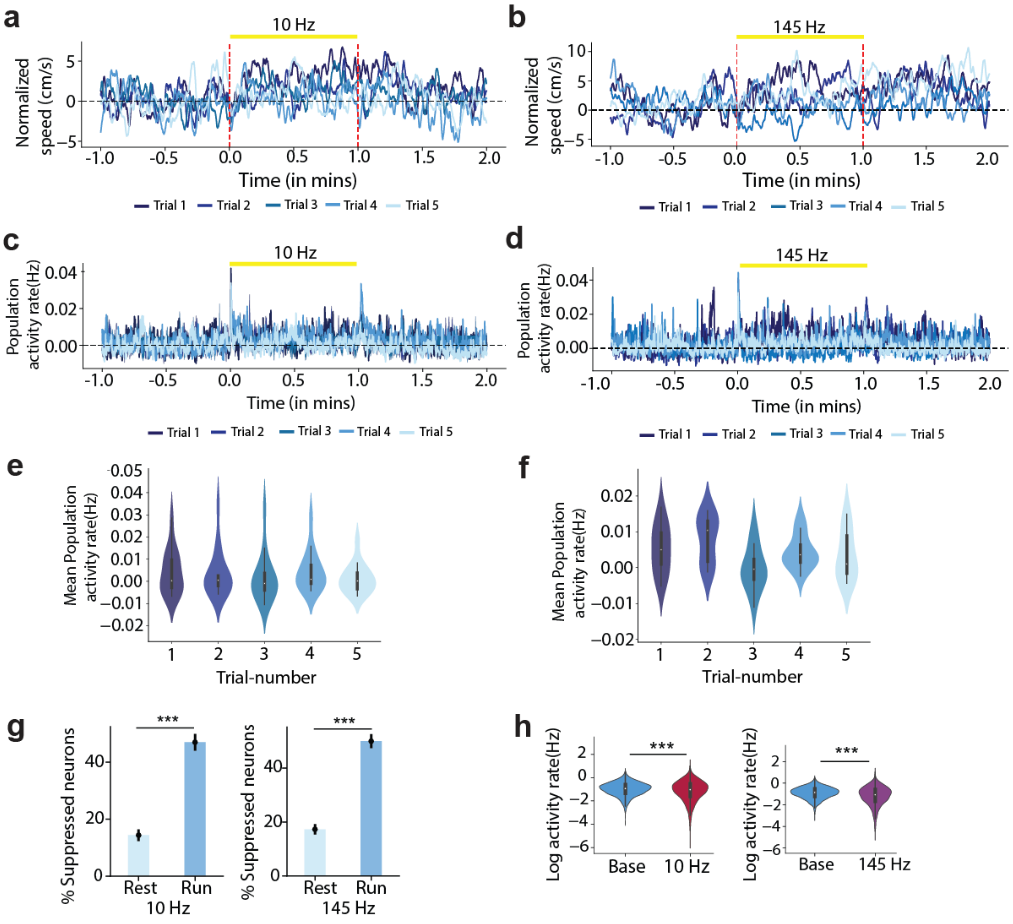


**Supplementary Figure 5**: **(a)** Population speed change (normalized to pre-stim period) across 5 trials of each 10 Hz stimulation session. Darker blue lines correspond to earlier trials. **(b)** same as (c), but for 145 Hz stimulation sessions. **(c,d)** Same as (a,b), but for population activity rate change across trials. **(e)** Mean population activity rate during stimulation across 5 trials of each 10 Hz session (Kruskal-Wallis test: p= 0.7, Post-hoc Dunn's test comparing first and fifth trial: p=1). **(f)** same as (e), but for 145 Hz stimulation sessions (Kruskal-Wallis test: p= 0.07, Post-hoc Dunn's test comparing first and fifth trial: p=1). **(g)** The fraction of movement responsive neurons that were suppressed by sensory stimulation during resting (light blue) and running (dark blue) across 10 Hz and 145 Hz stimulation sessions (Fisher's test, 10 Hz: $p=3.4e^{-64}$ (14.4% vs 47%), 145 Hz: $p=5.6e^{-78}$(17.3% vs 50%)). **(h)** Mean activity rate during baseline vs. stimulation at 10 Hz and 145 Hz (Mixed effect models, 10 Hz: $p_{approx}=3.7e^{-114}$, 145 Hz: $p_{approx}= 4.6e^{-158}$). Quantifications are visualized as violin plots with the outer shape representing the data kernel density and a box plot (box: interquartile range, whiskers: 1.5x interquartile range, white line: mean). *p < 0.05, **p < 0.01, ***p < 0.001

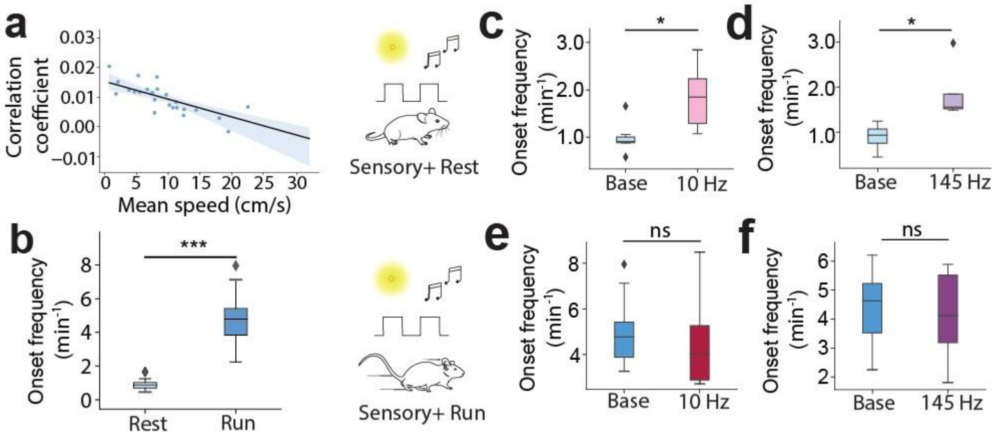


**Supplementary Figure 6: (a)** Median correlation coefficients across movement-responsive neurons vs. mean speed across the session. Each dot corresponds to a recording session. There is a significant linear relationship (linear regression, $R^2$=0.62, p=2.8$e^{-6}$, n=25 sessions) with shaded regions indicating the 95% confidence interval. **(b)** Session-wise mean pairwise activity rate during resting vs. running in baseline periods without stimulation (Wilcoxon signed rank test, p=2.7$e^{-5}$, n=25 sessions). **(c)** Session-wise mean pairwise activity rate across correlated neuron pairs during resting in baseline and for 10 Hz stimulation (Wilcoxon signed rank test, p=0.018, n=7 sessions) **(d)** Same as (**c**), but for 145 Hz stimulation (Wilcoxon signed rank test, p=0.028, n=6)**. (e)** Same as (**c**), but during running (Wilcoxon signed rank test, p=0.14). **(f)** Same as (**e**), but for 145 Hz stimulation (Wilcoxon signed rank test, p=0.07).

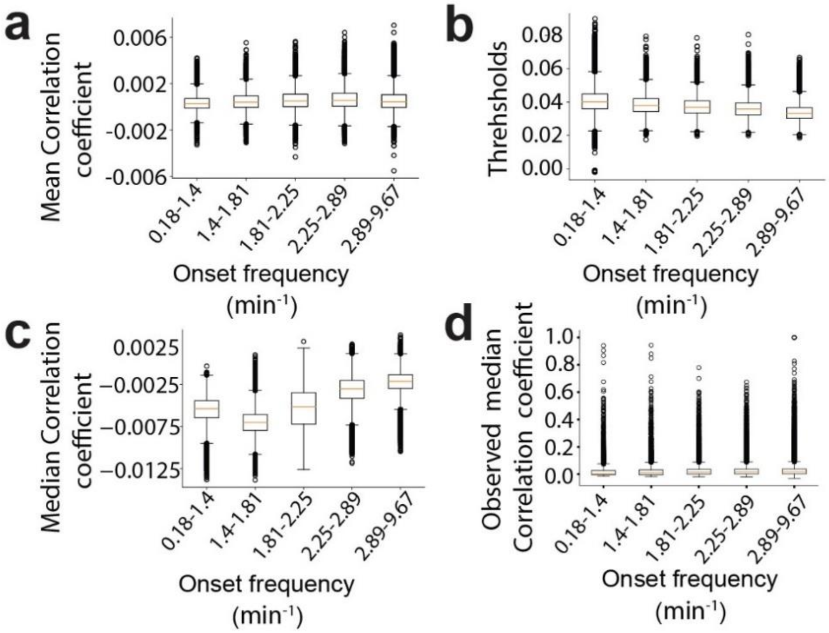


**Supplementary Figure 7: (a)** Mean correlation coefficients across shuffles versus calcium event onset rate (per min). **(b)** Thresholds (97.5$^{th}$ percentile) across shuffles versus calcium event onset rate. **(c)** same as a, but for median correlation coefficients across shuffles. **(d)** same as c, but for median correlation coefficients in the observed data.

### Tables

**Table 1: Sensory-responsive neurons during resting and running** (related to Fig. 5k)

| | **Sensory-responsive during resting** | **Number over total neurons recorded** | **Percentage** | **n** |
|---|---|---|---|---|
| **10 Hz** | Activated | 412/1813 | 22.72% | n=7 sessions, 5 mice |
| | Suppressed | 360/1813 | 19.86% | |
| | Activated + Suppressed | 772/1813 | 42.58% | |
| **145 Hz** | Activated | 562/2292 | 24.52% | n=6 sessions, 5 mice |
| | Suppressed | 482/2292 | 21.03% | |
| | Activated + Suppressed | 1044/2292 | 45.55% | |
| | **Sensory-responsive during running** | **Number over total neurons recorded** | **Percentage** | **n** |
| **10 Hz** | Activated | 1260/3155 | 39.94% | n=13 sessions, 9 mice |
| | Suppressed | 1214/3155 | 38.48% | |
| | Activated + Suppressed | 2474/3155 | 78.41% | |
| **145 Hz** | Activated | 909/2739 | 33.19% | n=13 sessions, 9 mice |
| | Suppressed | 1094/2739 | 39.94% | |
| | Activated + Suppressed | 2003/2739 | 73.13% | |

**Table 2: Fisher's test** (related to Fig. 5j)

| **Related to Fig. 5j** | | **Resting** | **Running** |
|---|---|---|---|
| 10Hz | Modulated | 712 (42.6%) | 2474 (78.4%) |
| | Non-modulated | 1041(57.4%) | 681 (21.6%) |
| 145Hz | Modulated | 1044 (45.5%) | 2003 (73.1%) |
| | Non-modulated | 1248 (54.5%) | 736 (26.9%) |

**Table 3: Overlap between sensory-responsive and movement-responsive neurons** (related to Fig. 6a)

| | **Category** | **Number over total neurons recorded** | **Percentage** |
|---|---|---|---|
| 10 Hz | Sensory-responsive during resting only | 75/1107 | 6.78% |
| | Sensory-responsive during running only | 557/1107 | 50.32% |
| | Both | 299/1107 | 27.01% |

| | | | |
|---|---|---|---|
| | None | 176/1107 | 15.9% |
| 145 Hz | Sensory-responsive during resting only | 135/1423 | 9.49% |
| | Sensory-responsive during running only | 601/1423 | 42.23% |
| | Both | 452/1423 | 31.76% |
| | None | 235/1423 | 16.52% |

**Table 4: Movement-responsive neurons that were also sensory-responsive** (related to Fig. 6b)

| | **Sensory-responsive during resting** | **Number over total neurons recorded** | **Percentage** | **n** |
|---|---|---|---|---|
| **10 Hz** | Activated | 215/1107 | 19.42% | n=7 sessions, 5 mice |
| | Suppressed | 159/1107 | 14.36% | |
| | Activated +Suppressed | 374/1107 | 33.79% | |
| **145 Hz** | Activated | 340/1423 | 23.89% | n=6 sessions, 5 mice |
| | Suppressed | 247/1423 | 17.36% | |
| | Activated +Suppressed | 587/1423 | 41.25% | |
| | **Sensory-responsive during running** | **Number over total neurons recorded** | **Percentage** | **n** |
| **10 Hz** | Activated | 336/1107 | 30.35% | n=7 sessions, 5 mice |
| | Suppressed | 520/1107 | 46.97% | |
| | Activated +Suppressed | 856/1107 | 77.33% | |
| **145 Hz** | Activated | 341/1423 | 23.96% | n=6 sessions, 5 mice |
| | Suppressed | 712/1423 | 50.04% | |
| | Activated +Suppressed | 1053/1423 | 74% | |

#### Table 5: Summary of all results

| **Category** | **Measure** | **Periods** | **10 Hz** | **145 Hz** |
|---|---|---|---|---|
| **LFP entrainment** | Power at the stimulation frequency (1 min before vs after onset) | Baseline vs. stim | ↑ | - |
| | LFP-stimulation pulse train phase locking | Baseline vs. stim | ↑ | ↑ |
| **Behavior** | % time in running | Baseline vs. stim | ↑ | ↑ |
| | Bout duration | Baseline vs. stim | ↑ | - |

| | | | | |
|---|---|---|---|---|
| | Motion onset frequency | Baseline vs. stim | ↓ | - |
| | Speed spectral delta power | Baseline vs. stim | ↑ | - |
| **LFP** | LFP-speed phase locking | True vs. shuffled - baseline and stim | ↑ | ↑ |
| | | Baseline vs. stim | ↑ | - |
| | Cross frequency coupling between speed-delta and LFP-beta | True vs. shuffled - baseline and stim | ↑ | ↑ |
| | | Baseline vs. stim | ↑ | - |
| | Cross frequency coupling between speed-delta and LFP-gamma | True vs. shuffled | ↑ | ↑ |
| | | Baseline vs. stim | - | - |
| **Calcium responses** | Event rate (Mixed effect models – Movement-responsive neurons) | Resting - stim vs. baseline | - | - |
| | | Running - stim vs. baseline | ↓ | ↓ |
| | Cross correlation - lag | Baseline vs. stim | - | - |
| | Cross correlation – correlation coefficient | Baseline vs. stim | - | - |
| **Ca-movement** | $Ca^{2+}$ vs. delta-component of speed PLV | True vs. shuffled- baseline and stim | ↑ | ↑ |
| | | Baseline vs. stim | - | - |
| | Preferred phase (2-4 Hz) | Baseline vs. stim | - | - |
| **Ca-LFP** | $Ca^{2+}$ vs. delta component of LFP PLV | True vs. shuffled- baseline and stim | ↑ | ↑ |
| | | Baseline vs. stim | - | - |
| | Preferred phase (2-4 Hz) | Baseline vs. stim | - | - |
| **Pair-wise Correlation** | Effect of Locomotion | % Correlated pairs | ↑ | ↑ |
| | | Correlation coefficient across correlated pairs | ↓ | ↓ |
| | | Corr. Coeff. across all pairs of movement-responsive neurons | ↓ | ↓ |
| | | Event onset frequency | ↑ | ↑ |
| | Effect of stimulation during resting | % Correlated pairs | ↓ | ↓ |
| | | Correlation coefficient across correlated pairs | ↑ | ↑ |
| | | Event onset frequency | ↑ | ↑ |
| | Effect of stimulation during running | % Correlated pairs | ↓ | ↓ |
| | | Correlation coefficient across correlated pairs | ↑ | ↑ |
| | | Event onset frequency | - | - |
